# Linking transcriptome to cell behavior in real time uncovers molecular fate-asymmetry and a quiescence cycle among adult neural stem cells

**DOI:** 10.64898/2026.07.05.736547

**Authors:** Tanya Foley, Isabelle Foucher, Gaëlle Letort, Jules Samaran, Marion Dussauge, David Morizet, Nicolas Dray, Laura Cantini, Laure Bally-Cuif

## Abstract

Adult neural stem cells (NSCs) are transcriptionally heterogeneous, yet the relationship between molecular heterogeneities and individual NSC trajectories remains unclear. Here, we link transcriptional identities with cellular features and measures of real time to identify molecular trajectories hidden within transcriptomic space. Among self-renewing NSCs, we resolve single-cell molecular transitions associated with fate asymmetry at division. We also show that individual self-renewing NSCs progressively transition from molecular states of deep to shallow quiescence during prolonged quiescence phases. Together, this work reveals individual NSC trajectories within transcriptomic space during fate decisions and state transitions. In particular, it highlights the existence of a transcriptionally encoded quiescence cycle followed by adult NSCs, independent of lineage progression, that balances division with cell growth to sustain self-renewal over time.

## INTRODUCTION

To balance neurogenesis with long-term maintenance, adult neural stem cells (NSCs) in the vertebrate brain alternate between fate-indicating divisions and quiescence phases, during which they experience prolonged yet reversible cell cycle arrest^1–3^. Among NSC, quiescent phases are essential to maintaining stemness, with continued proliferation leading to NSC exhaustion^4^. Fate-indicating divisions following quiescence exit are either asymmetric to permit individual self-renewal and the production of progeny fated to neurogenesis, or symmetric to amplify the NSC or progenitor pools^1,2,5–7^. The molecular mechanisms that govern NSC fate and state decisions remain incompletely understood. Closing this knowledge gap necessitates linking cellular behaviors, measured in real time and space, with changes in individual NSC molecular identity, or transcriptional trajectories.

NSCs tracked in real time by live imaging and clonal analyses has revealed heterogeneities in behavior, including proliferation frequencies, division modes, and quiescence durations^1–3,5,8,9^. Transcriptomic assays have similarly identified single-cell NSC heterogeneities, corresponding to differences in regional identity, lineage progression, and quiescence depth^10–20^. How transcriptional identity relates to individual cell behavior over time, however, remains unaddressed, as this requires experimental measures of real time that are difficult to combine with omics-based approaches. While methods for pseudotime trajectory inference have been developed, such methods identify the most probable cell state transitions at population-level, relying on the assumption that transitions are continuous and likely to occur between most closely related states^21^. As such, these approaches are best-suited for continuous transitions over short timescales^22–25^, yielding trajectories that are difficult to interpret when applied to populations like homeostatic adult stem cell systems, in which dynamics are slow with state transitions that are complex and discontinous^26^. Longitudinal time-series comparisons in developmental systems can overcome these limitations^27^, however adult NSCs are subjected to population homeostasis by which a constant representation of substates is maintained over time, despite changes to individual cell identities^28^. Thus, unlike in developmental systems, state transitions cannot be inferred through a time-series comparison. Such challenges obscure our understanding of molecular transitions associated with lineage progression at division, as well as possible cell-state transitions within quiescence. Quiescence is an actively maintained and transcriptionally a heterogeneous state^12,29^, with quiescence depth, or resistance to activation, graded across NSC populations^16–18,30^. In previous work, short-term pharmacological perturbation of pro-quiescence Notch signaling produced a graded activation response by NSCs, suggestive of different baseline quiescence depths^17^. Whether quiescence heterogeneity reflects distinct NSC subpopulations, or rather asynchronous transitions between quiescent substates, remains unclear.

To connect NSC behavior and transcriptome, we leveraged here the adult zebrafish (*Danio rerio*) dorsal telencephalon (pallium), where NSC behavior has been resolved in real time at population scale^1,2^. By intravital imaging, individual NSC behaviors like proliferation, lineage progression, and fate decisions were previously characterized over time, identifying a population a self-renewing NSCs biased toward asymmetric divisions and subjected to quiescence phase estimated to persist for several months^1,2,5,6,9^. We reasoned that these real-time measurements of NSC state and fate transitions could serve as a reference frame for interpreting single-cell transcriptional dynamics across quiescence and at division. To this end, we adapted multiplexed single-molecule RNA fluorescence in situ hybridization (smFISH) to the whole mount pallium to examine transcriptional heterogeneities in the context of cell morphology and kin relationships. To elucidate transitions between molecular NSC states, we combined transcriptional identity with measures of real time via BrdU time-stamping. With this, we distinguish between molecular trajectories associated with fate progression and those underlying transitions within quiescence. We show that transcriptionally encoded fate asymmetry at division immediately partitions daughter cells into distinct transcriptional pools. We also show that upon quiescence re-entry, self-renewing daughter NSCs first experience a molecular state of deep quiescence that is resolved over extended quiescence phases as NSCs transcriptionally progress toward shallow quiescence. Together, through a multi-modal approach connecting transcriptional identity with cell behavior, we uncover single-cell molecular trajectories, both at division and within quiescence, that underly constitutive adult neurogenesis.

## RESULTS

### A minimal transcriptomic code assigns subcluster identities to pallial neural stem and progenitor cells in situ

Single-cell RNA-sequencing (scRNAseq) was previously performed on cells from the dorsal telencephalon (pallium) of Tg(*sox2:gfp*) adult zebrafish at 3-months post-fertilization (mpf)^20^. Among profiled cells, quiescent neural stem and progenitor cells (NSPCs) were identified by the expression of astroglial markers in the absence of *pcna*^20^ (**Figures 1a and 1b**). Subsequent subclustering identified seven quiescent (q) subclusters (q1-q7) and three proliferating clusters (p1-p3), with subclusters q1-q5 expected to contain quiescent pallial NSPCs (**Figures 1c and 1d**)^20^. These NSPCs are highly accessible for imaging in situ, forming a monolayer at the ventricular surface of the pallium with cell bodies that span the first 10-15µm in depth. To examine subclusters q1-q5 within their intact niche, we defined a minimal transcriptomic code composed of 12 genes that, when measured together, distinguished between q1-q5 (**Figures 1e and 1f**). To detect these genes in situ, we adapted single molecule RNA-fluorescence in situ hybridization (smFISH) for whole mount pallia. This was combined with fluorescence immunohistochemistry (IHC) against Zonula occludens-1 (Zo1), marking tight junctions delineating apical cell boundaries, and the nuclear NSPC marker Sox2^1,31^. Probes for the 12-gene transcriptomic code were multiplexed in situ, revealed three-by-three across four iterative rounds of imaging to capture gene expression among NSPCs in the anterior and medial pallial regions (Da/Dm) (**Figures 1g, 1h, and S1a**)^9^.

**Figure 1.**
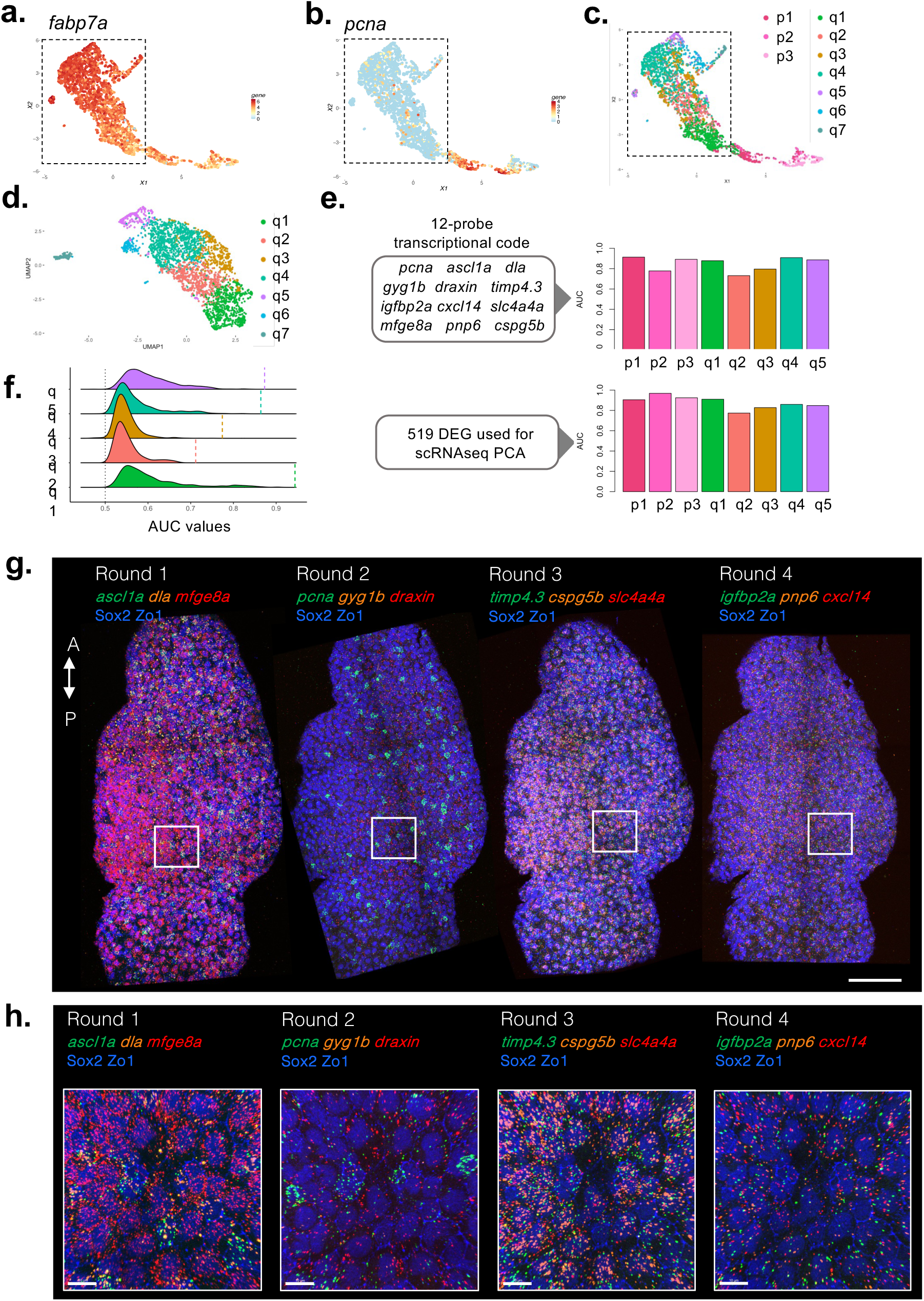
A minimal “Transcriptomic Code” based on scRNAseq analyses can be revealed in NSPCs *in situ*. **a, b.** *fabp7a* (a) and *pcna* (b) expression across the NSPC population visualized by UMAP with quiescent progenitors (*pcna^neg^*) indicated by a dashed box^20^. **c.** UMAP of NSPCs color-coded by sub-cluster identity^20^. **d.** UMAP of quiescent NSPCs only color-coded by subcluster^20^. **e.** Predicted accuracy of substate identification using the 12-gene transcriptomic code compared to all 519 variable genes applied to PCA analysis of scRNA-seq data as determined by area under the receiver-operator curves (AUC). **f.** Monte Carlo simulation reinforces gene selection, with the distribution of AUC values from 1000 receiver-operator simulations using 12 transcripts randomly selected among those detected within NSPCs by scRNA-seq far from values obtained using the 12-gene transcriptomic code, indicated by dashed lines. **g.** 3D reconstruction of a pallial hemisphere (dorsal view) following four iterative rounds of multiplexed smFISH (green, orange, red) with immunostaining (blue). A, anterior; P, posterior. Scale bar, 100µm. **h.** Magnified pallial regions indicated by the boxed areas in (f) showing mRNA puncta. Scale bars, 10μm.

To quantify gene expression in whole-mount tissue, we developed image-analysis tools, including *Multireg*, to combine iterative confocal images, and *FishFeats,* to measure gene expression in situ, along with apical cell area using segmented Zo1 staining (**Figures S1b-e**)^32^. We observed patterns of gene expression across a transcriptional continuum highly concordant with those by scRNAseq (**Figure 2a**)^20,32^. By both modalities, *pnp6* and *cspg5b* were enriched at one end of the continuum, overlapping in expression with the NSC marker *mfge8a*^33^, *slc4a4a* and *cxcl14*. *timp4.3* expression was coincident with that of *igfbp2a* and with *draxin*, required for adult neurogenesis in the mouse hippocampus^34,35^. *gyg1b* was expressed at low levels, in contrast with *dla* and *ascl1a,* both being highly expressed at one end of the continuum and partially overlapping with the cell cycle marker *pcna*. These gene expression patterns, summarized by correlation matrix (**Figure S1f**), were reproduced in a second biological replicate (data not shown). To assess non-specific signal, a separate pallium was processed in parallel in which control probes were applied targeting bacterial L-2,3-dihydrodpicolinate reductase transcript (*dapB),* corresponding to each of the 12 probes within the transcriptomic code (data not shown). As considerably fewer puncta per cell were detected compared to “true” signal, no corrective factor was applied to in situ gene expression values.

**Figure 2.**
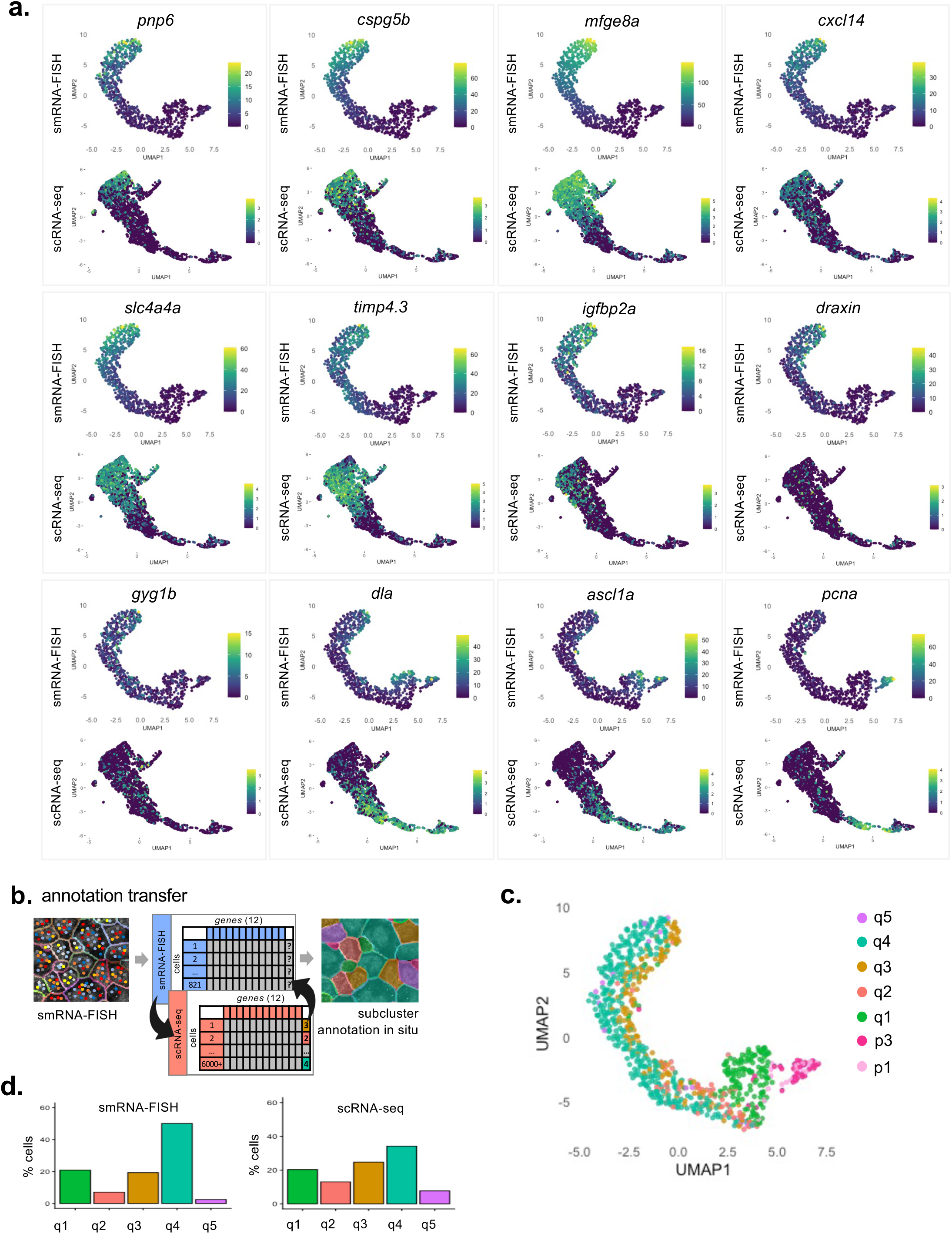
Concordance in gene expression patterns between single-cell transcriptomic modalities allows for identification of q-subclusters q1-q5 in the whole mount pallium. **a.** UMAPs for raw gene expression values as detected by multiplexed smFISH for the 12-gene transcriptomic code compared to normalized gene expression detected by scRNAseq^20^. **b.** Subcluster annotation transfer from scRNAseq to cells for which gene expression is measured by smFISH. **c.** UMAP showing q-subcluster annotations transferred from scRNAseq onto cells measured in situ. **d.** Percent cells among each q-subcluster identity for cells measured in situ following annotation transfer (left) compared to direct subclustering from scRNAseq (right).

To transfer scRNAseq subcluster identities onto NSPCs measured in situ, smFISH cells were projected into scRNAseq transcriptomic space using expression values for the 12 genes common to both modalities, and the most common q-subcluster identity among the 20 closest scRNAseq neighbors was assigned to cells assessed in situ (see methods) (**Figure 2b**). 817 of the 821 cells considered were successfully annotated to subcluster identities q1-q5, p1, and p2 (**Figure 2c**). Cells with identities q1 and q2 were found at one extreme of the transcriptional continuum, proximal to cycling cells, with q3, q4, and q5 enriched at the opposing end. The relative proportions of smFISH cells annotated as q1-q5 was comparable to that by direct scRNAseq subclustering (**Figure 2d**)^20^. Together, the patterns of gene expression conserved across both modalities, as well as the recovery of all expected subcluster identities following annotation transfer, highlight the robustness of our smFISH method and the selection of target genes. Further, these data demonstrate that NSPC heterogeneities typically encoded by the entire transcriptome are sufficiently captured using only 12 differentially expressed genes measured at single-molecule resolution in situ.

### Multimodal analysis in situ positions self-renewing NSCs within a transcriptional continuum dominated by lineage progression

We previously identified a hierarchy of NSPC subpopulations sustaining stemness maintenance, in which upstream “reservoir” NSCs divide asymmetrically to self-renew and fuel a downstream “operational” pool that is more proliferative and further committed to neurogenesis^1,2^. To explore transcriptional heterogeneities in situ using this framework, we performed hierarchical clustering on smFISH data (**Figures 3a and S2a**). Two broad clusters were identified, clusters 1^12^ and 2^12^ (with superscript referring to the number of genes used for clustering), with individual cells assigned cluster stability scores (see methods) (**Figure S2b**).

**Figure 3.**
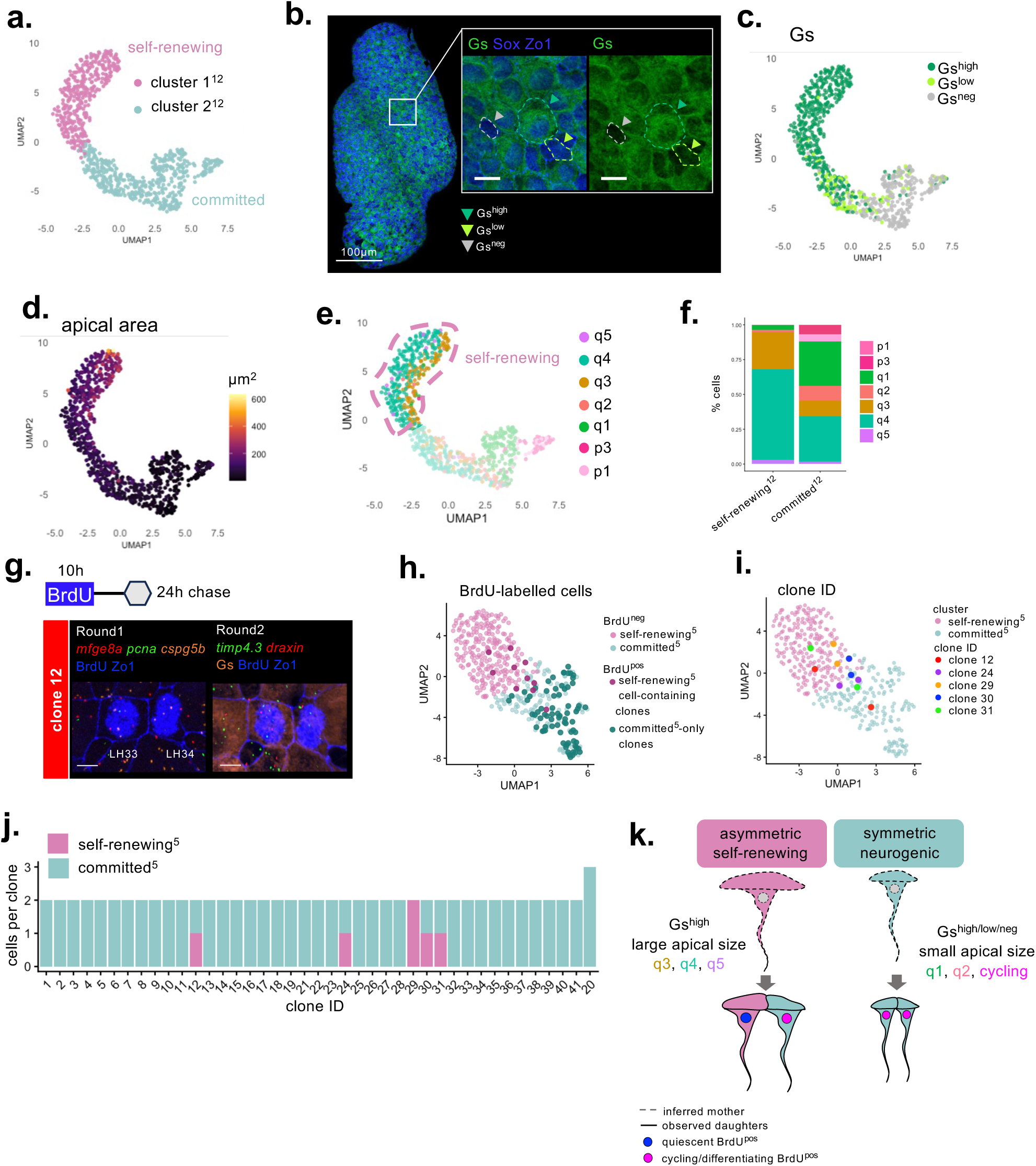
Gene expression in situ combined with histological features and BrdU tracking of division modes allows for self-renewing NSCs to be transcriptionally identified. **a.** NSPC clusters identified by hierarchical clustering of smFISH data, with clusters 1^12^ and 2^12^ containing 355 and 462 cells, respectively. **b.** Immunostaining for Gs and Zo1 in a whole mount pallium following 12-gene smFISH. Cells were classified as Gs^high^, Gs^low^, or Gs^neg^. Dashed lines indicate representative cells from each category. Scale bar, 10µm. **c-e.** UMAP representations of NSPCs measured by smFISH and color-coded by Gs intensity (c), apical area (d), and q-subcluster annotation with the self-renewing^12^ cluster outlined by dashed lines (e). **f.** Percent cells assigned to scRNAseq subcluster identities within the self-renewing^12^ and committed^12^ in situ clusters. **g.** Representative BrdU-doublet (clone 12 in panel (i)) among those 24h post-division assayed for *mfge8a*, *pcna*, *cspg5b*, *timp4.3*, and *draxin* expression with immunostaining for Zo1, BrdU, and Gs. Scale bar, 5µm. **h.** 435 pallial NSPCs measured for *mfge8a*, *pcna*, *cspg5b*, *timp4.3*, and *draxin* expression treated with a 10-hour BrdU pulse followed by a 24-hour chase. Cluster identities and BrdU-labelled cells are indicated on the UMAP by color-coding. **i.** Sister cells within self-renewing^8^ cell-containing doublets 24h post-labelling indicated by color-coding. **j.** Number of cells within BrdU^pos^ clones 24h post-division and their cluster identities. **k.** Summarized schematic representation of division modes and cellular features tracked by the 10-hour BrdU pulse followed by a 24-hour chase.

To determine the biological significance of clusters 1^12^ and 2^12^, we examined the distribution of cellular features associated with lineage progression (glutamine synthetase – Gs – intensity and apical size) across transcriptional space^1,36^. For this, we complemented multiplexed smFISH with IHC for the NSC marker Gs, captured in a fifth round of imaging^2,36^. Using *FishFeats*, we manually classified cells as Gs^high,^ Gs^low^, or Gs^neg^ (negative), distinguishing NSCs (Gs^pos^) from committed neural progenitors (NPs) (Gs^neg^) (**Figure 3b**). We found that transcriptionally, Gs^high^ NSCs were enriched at one end of the continuum, opposite Gs^neg^ NPs (**Figure 3c**). Assessing apical area similarly revealed graded variability in apical size across the continuum, with larger cells being Gs^high^ and smaller cells mostly Gs^neg^, consistent with previous work showing an inverse relationship between apical size and potential for self-renewal^1^.

Next, we considered how these features were distributed across clusters 1^12^ and 2^12^. We found that cluster 1^12^ cells were almost exclusively Gs^high^, while cluster 2^12^ varied in Gs intensity yet contained all Gs^neg^ NPs (**Figure S2c**). Cluster 1^12^ also contained cells large in mean apical size (168 ± 74µm^2^) relative to those in cluster 2^12^ (54 ± 40µm^2^) (**Figures 3d and S2d**). Together, these data confirm that cluster 1^12^ is enriched in NSCs, while cluster 2^12^ includes further committed cells, with lineage progression being the main driver of heterogeneity along the major axis of the NSPC continuum. Based on differences in stemness potential, we hypothesized that clusters 1^12^ and 2^12^ might represent the reservoir and operational NSPC pools^1,2^. To test this, we used pallia isolated from Tg(*dla:gfp*)^37^ adults, in which a GFP reporter driven by the expression of the Notch ligand *deltaA* (*dla*) marks operational cells with high GFP levels, while reservoir NSCs remain null ^1^ to weak (unpublished observations)^1^. Comparing Gs intensities, apical area, and *mfge8a* expression between clusters 1^12^ and 2^12^, and between reservoir and operational cells, demonstrated that cluster 1^12^ was enriched in reservoir NSCs while cluster 2^12^ approximated the operational pool (**Figures S2e-k**). For simplicity, these clusters are referred to as “self-renewing” (cluster 1^12^) and “committed” (cluster 2^12^) hereafter.

Finally, to reconcile scRNAseq-derived subclusters with self-renewing^12^ and committed^12^ cluster identities, we examined the distribution of q1-q5 and p1-p3 annotations across these clusters. We found the self-renewing^12^ cluster to be nearly exclusively composed of cells with q3, q4, and q5 annotations (**Figures 3e and 3f**). In contrast, a large percentage of cells within the committed^12^ cluster were annotated as q1, p1, and p3. Thus, in addition to positioning functionally distinct NSPC subpopulations within transcriptional space, these findings indicate that scRNAseq subcluster identities q3, q4, and q5 represent transcriptional substates specific to self-renewing NSCs during quiescence.

### Transcriptional asymmetry at division separates self-renewing and committed NSPC populations

Using intravital imaging, we previously showed that operational cells were generated from self-renewing reservoir NSCs through fate-asymmetry at division^1^. Now, with access to transcriptomic identities in situ, we examined whether fate asymmetry at division could be detected at the transcriptional level, with daughter cells partitioned between different transcriptomic subspaces. To this end, adult zebrafish were treated with a 10-hour bromodeoxyuridine (BrdU) pulse, labelling cells in S-phase. Labelled NSPCs were identified by IHC 24 hours post-division, with no BrdU singlets observed, indicating all labelled cells completed mitosis during the chase without G2 arrest. 41 BrdU-labelled clones were identified, consisting of 40 doublets and 1 triplet (**Figure 3g)**. These cells were assayed for *cspg5b*, *mfge8a*, *timp4.3*, *draxin*, and *pcna* expression by smFISH, along with 352 BrdU^neg^ cells within the same hemisphere (435 cells total), generating a transcriptional continuum along which BrdU^pos^ cells could be visualized (**Figures 3h and S3a**). Hierarchical clustering again separated cells into two broad clusters, clusters 1^5^ and 2^5^, largely approximating the self-renewing^12^ and committed^12^ populations based on Gs intensities, apical cell sizes, and patterns of gene expression (**Figures 3h and S3a-d**). Among the 41 BrdU-labelled clones, only 5 contained at least one self-renewing^5^ daughter cell, with all other clones comprised of only committed^5^ progeny. Among these 5 clones, 4 were asymmetrically partitioned between the self-renewing^5^ and committed^5^ clusters (**Figures 3i and 3j**). Exceptionally, both cells within clone 29 (RH128 and RH129) were self-renewing^5^, with RH128 relatively low in its cluster stability score (**Figure S3d**).

Together, these results reveal the existence of two division modes at the transcriptional level: rare asymmetric divisions by which two daughter cells are partitioned between the self-renewing^5^ and committed^5^ clusters, and more frequent amplifying divisions resulting in only committed^5^ progeny. While the transcriptional identity of mother cells pre-division cannot be assessed by our approach, previous characterizations of NSC division modes allow us to infer that the asymmetric divisions tracked here originate from self-renewing^5^ mother cells^1^. As such, these data demonstrate that asymmetric divisions by self-renewing^5^ mother cells generate a transcriptional boundary between the self-renewing^5^ and committed^5^ clusters. Further, that one of two daughters are found within the self-renewing^5^ cluster suggests that such daughters are equivalent to their mothers in stemness potential, and to quiescent self-renewing^8^ cells that remain unlabelled by division (**Figure 3k**).

### Long-term BrdU time-stamping links transcriptional identity with cell morphology, lineage relationships, and fate in real time

To interrogate the relationship between transcriptional identity and cell behavior over time, we applied BrdU time-stamping. For this, adult zebrafish were treated with a 10h BrdU-pulse followed by increasing chase times post-labelling, at which transcriptional identities among BrdU-labelled NSPCs were assessed. Pallia were collected 4-days (4d), 8d, 14d, and 28d after the pulse, revealing sparse BrdU-labelling throughout Da/Dm (**Figure 4a**), as before. The short pulse, being shorter than the estimated cell cycle length^38–41^, labelled only one division within a lineage, allowing for subsequent amplifying divisions to be tracked by inheritance of the BrdU label. This, combined with the absence of both lateral migration and cell death among dorsal NSPCs and their progeny, meant that transcriptional identity could be connected with lineage relationships over time^2,5,7,9,42^. We assayed gene expression for 8 targets within the 12-gene transcriptomic code (*pcna*, *ascl1a*, *dla*, *gyg1b*, *draxin*, *timp4.3*, *mfge8a*, and *cspg5b*, selected for their complementary patterns of expression and combined accuracy in cell state identification) (**Figures 4b, S4a and S4b**). Gs intensities and apical cell sizes were also assessed by IHC (**Figures 4b, 4e and 4f**). For all chase times, BrdU^pos^ cells with an apical surface within the NSPC layer were quantified across both pallial hemispheres, along with enough BrdU^neg^ cells to generate a transcriptional continuum onto which BrdU^pos^ cells could be positioned (**Figures 4c-g**). In total, 5106 cells from BrdU pulse-chase replicates and the initial dataset (**Figure 1f**, “no BrdU”) were integrated, producing the expected patterns of gene expression across a transcriptional continuum (**Figure S4c**). Hierarchical clustering again identified two major clusters, clusters 1^8^ and 2^8^ (**Figures 4d, S4d and S4e**), once again approximating self-renewing and committed NSPC populations (**Figures 4d-f, S4c-h**).

**Figure 4.**
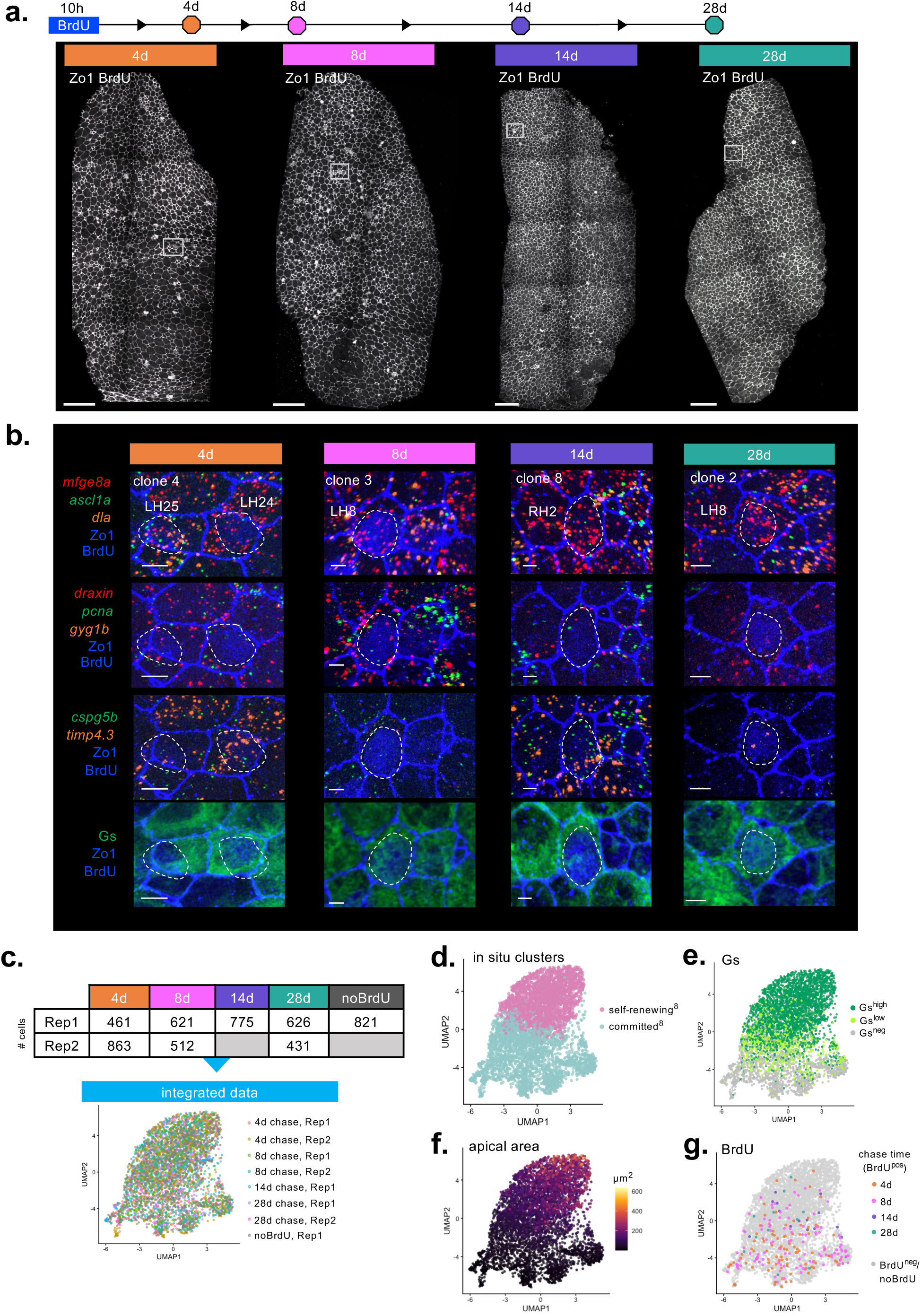
Validation of BrdU time-stamping to link cell behavior with transcriptional identity over time in situ. **a.** 3D reconstruction of pallial hemispheres treated with a 10-hour BrdU pulse followed by increasing chase times post-labelling. Boxed regions highlight BrdU^pos^ cells magnified in (b). Scale bars, 70µm. **b.** Representative BrdU^pos^ cells at each chase time for which gene expression was measured by smFISH with immunostaining for Zo1, BrdU, and Gs. BrdU^pos^ cell IDs are indicated (Round 1) with dashed lines surrounding BrdU^pos^ nuclei. Scale bars, 5µm. **c.** The number of NSPCs from 8 integrated smFISH datasets visualized by UMAP. **d.** Self-renewing^8^ and committed^8^ clusters within the NSPC UMAP following hierarchical clustering of integrated smFISH data. **e-g.** Gs intensity (d), apical surface size (e), and the position of BrdU-labelled cells at each chase time (f) across the integrated NSPC UMAP.

### Real-time single-cell trajectories reveal that transcriptional asymmetry post-division signs fate asymmetry

Using these integrated data, we first aimed to determine whether transcriptional asymmetry 24h post-division (**Figure 3g-k**) signs fate asymmetry. We compared transcriptional identities among BrdU-labelled cells within clones at increasing chase times, with each clone representing an individual cell lineage (**Figure 5, S5, and S6**). The number of cells maintaining an apical surface was quantified for all clones at each chase time (**Figures 5a-e**). Clones possessed between 1-3 cells with apical surfaces across chase times, with only one clone at 4d post-division containing 5 such cells (**Figure 5b**). Given that all labelled cells transitioned from S-phase to mitosis within 24h (**Figure 3h**), clones in which only a single apical domain-containing cell were observed necessarily had a sister at the time of labelling. These sisters, having delaminated during the chase (along with any progeny), were not considered for gene expression analysis.

**Figure 5.**
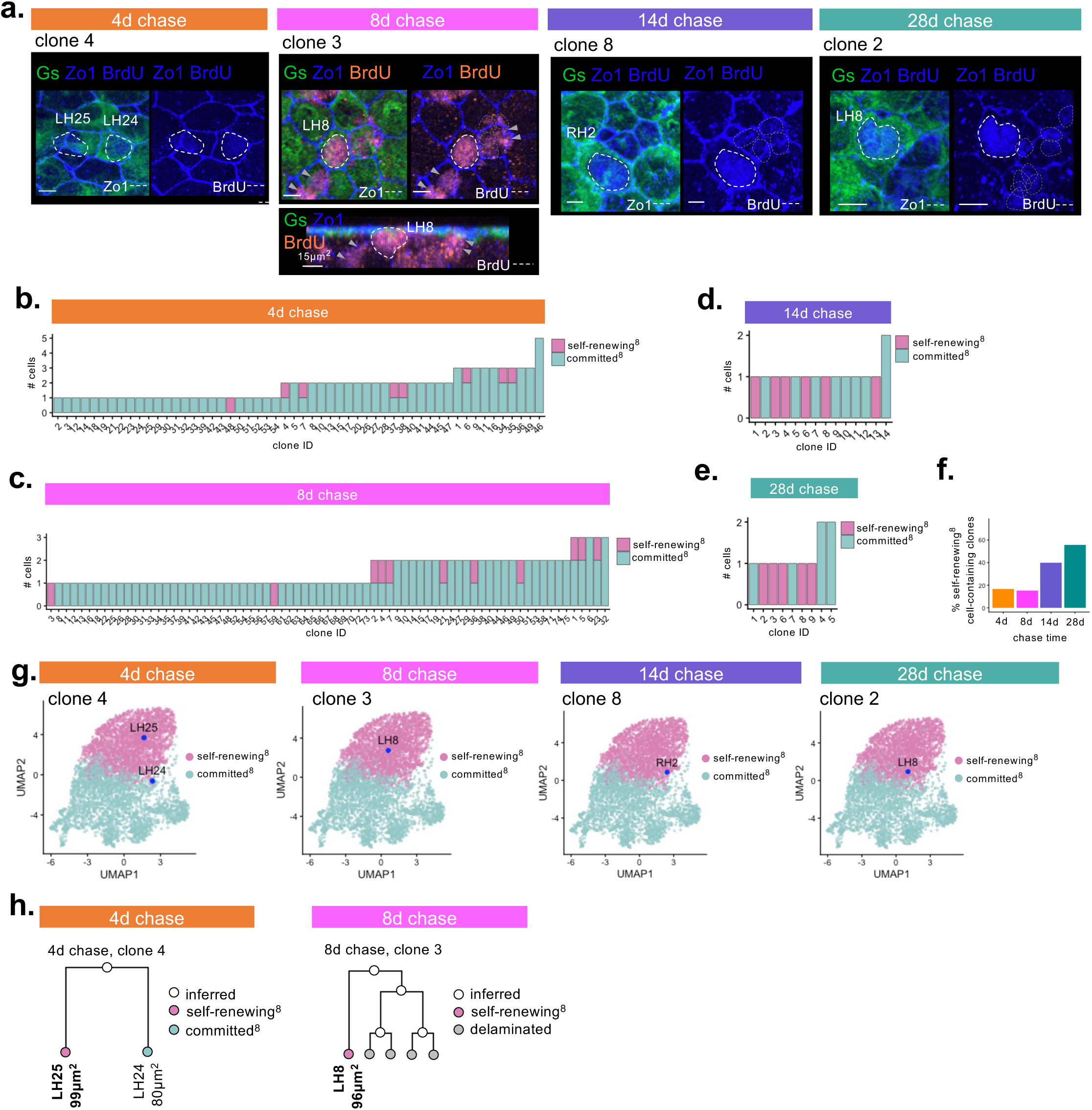
Transcriptional asymmetry between progeny of self-renewing NSCs resolves into fate asymmetry at increasing chase times. **a.** 3D reconstructions of pallial hemispheres treated with a 10-hour BrdU pulse at the indicated chase times in which immunostaining for BrdU, Zo1 and Gs are shown. White dashed lines indicate nuclear BrdU among cells maintaining an apical surface. Grey dashed lines indicate BrdU^pos^ nuclei belonging to delaminated cells within the clone of interest. Grey arrowheads indicate BrdU^pos^ nuclei in the vicinity of, but apart from, clones of interest. BrdU^pos^ clones at 8d post-division were examined in optical cross-section (lower panel). Scale bars, 5µm, unless otherwise specified. **b-e.** Number of cells per BrdU^pos^ clone maintaining an apical surface at 4- (a), 8- (b), 14- (c), and 28-days (d) post-division. Cells within clones are color-coded by cluster identity. **f.** Percentage of self-renewing^8^ cell-containing clones at increasing chase times post-labelling. **g.** Position of cells from individual BrdU-labelled clones within the integrated UMAP. **h.** Inferred lineage trees for clones at 4d and 8d post-labelling. Cell ID and apical cell size are indicated for cells maintaining an apical surface.

At 4 and 8d post-division, most BrdU-labelled cells were committed^8^ (46/54 and 62/73 at 4 and 8d, respectively) (**Figures 5b and 5c**). Among rare self-renewing^8^ cell-containing clones, we observed only one self-renewing^8^ cell per clone (**Figures 5b-e, 5g, and S6**). This is consistent with fate asymmetry at division, by which one daughter cell (being self-renewing) returns to quiescence. At later chase times, the total number of BrdU-labelled clones at the pallial surface was reduced, with 6/14 such clones containing a single self-renewing^8^ cell 14d post-labelling, compared to 5/9 at 28d (**Figures 5d, 5e**). Despite the decrease in total clone number, the proportion of clones containing self-renewing^8^ cells increased over time (**Figure 5f**), concomitant with an accumulation of BrdU^pos^ nuclei below the pallial germinal layer (**Figures 5a and S5**).^5,7,9,43,44^ Together, persistence of self-renewing^8^ cells within the pallial germinal layer, along with the loss of committed^8^-only clones, is consistent with delamination into the underlying parenchyma by BrdU-labelled committed^8^ progeny undergoing terminal differentiation during the chase^5,7,9,43,44^.

Previous work showed that divisions by NSPCs produce daughter cells similar in apical size, each being approximately half the size of the mother^1^. This is visible within clone 4 at 4d post-division with LH24 and LH25 being 99µm^2^ and 80µm^2^ in apical size, respectively (**Figures 5a** (4d chase) **and 5h**). We inferred lineage trees for self-renewing^8^ cell-containing clones based on the total number of BrdU^pos^ nuclei, their relative intensities, and the apical area measured among cells within the clone (**Figures 5h, S5b and S5d**). Clone 35 at 4d post-division, for example, contained three apical domain-containing cells: RH6, RH7, RH8 (**Figures S5a and S5b** (clone 35)). Given its transcriptional identity, RH8 is assumed to have returned to quiescence post-division, consistent with its large apical size. The apical sizes of RH6 and RH7, however, being approximately half that of RH8 and comparable to one another, suggest these cells are likely to be sisters, produced by a second amplifying division following that by which RH8 was labelled. This interpretation is further consistent with the transcriptional identities of RH6 and RH7, both being within the committed^8^ cluster (**Figures S5b and S6a** (clone 35)).

Taken together, these observations demonstrate that by tracking single-cell transcriptional identities within individual lineages, state transitions can be linked with cell behavior over time. As only one self-renewing^8^ cell was identified per self-renewing^8^ cell-containing clone across all chase times, systematically large in apical area, we show that self-renewing^8^ cells maintain quiescence post-division over a period of at least 28d. In contrast, we find that committed^8^ progeny undergo additional amplifying divisions, becoming progressively lost from the NSPC layer over time as they engage in differentiation. Finally, we reveal differences in proliferation frequency, division mode, and capacity for self-renewal between clusters 1^8^ and 2^8^, consistent with reservoir and operational identities, respectively.

### Among self-renewing NSCs, transcriptional heterogeneity reflects a continuum of variable quiescence depths

Among subclusters q1-q5, scRNAseq revealed differential sensitivities to transient Notch blockade^17^. While this observation suggested that q-subcluster identities likely correspond to different quiescence depths, such differences could not be specifically attributed to self-renewing NSCs, nor could individual NSC trajectories within quiescence be reconstructed. To address these questions, we restricted our analysis of transcriptional heterogeneities to quiescent self-renewing^8^ cells, identifiable in situ. Among these cells, we re-integrated those with cluster stability scores ≥ 0.95 (**Figure 6a**), with the resulting subset largely restricted to Gs^high^ cells (**Figures 6b and S7a**).

**Figure 6.**
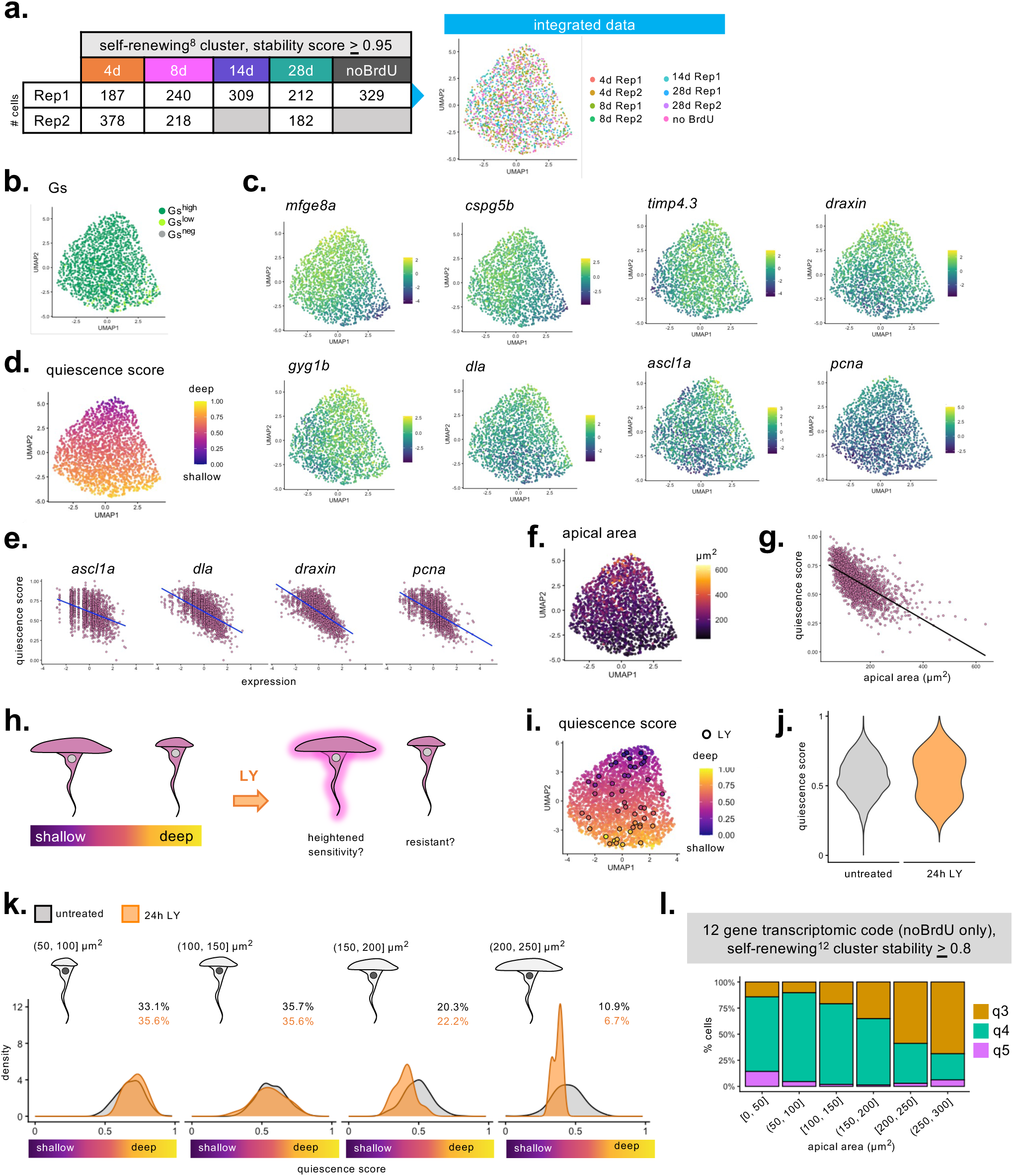
A continuum of variable quiescence depths underlies transcriptional heterogeneity among self-renewing NSCs. **a.** The number of self-renewing^8^ cells from each of the integrated smFISH datasets (left) and their visualization by UMAP (right). **b-d.** Gs intensities (b), gene expression (c), and quiescence scores across the self-renewing^8^ continuum visualized by UMAP. **e.** Quiescence score as a function of *ascl1a*, *dla*, *draxin*, and *pcna* expression within the self-renewing^8^ continuum. **f.** Apical area among self-renewing^8^ cells visualized by UMAP. **g.** Quiescence score as a function of apical cell area within the self-renewing cluster^8^, with a trendline derived by linear regression. **h.** Expected responses to short-term Notch inhibition based on apical size, with large self-renewing^8^ cells in shallow quiescence exhibiting a heightened response to Notch inhibition. **i.** Quiescence scores for 2522 untreated and 47 LY-treated (black outline) cells within the self-renewing^8^ cluster visualized by UMAP. **j.** Distribution of quiescence scores among untreated and LY-treated cells. **k.** Kernel density plots showing the distribution of untreated and LY-treated self-renewing^8^ NSCs across the range of quiescence scores with cells binned by apical size (50µm^2^ bins). Cells above 250µm^2^ were considered outliers based on the interquartile range. The comparable percentages of cells from each condition distributed across apical size bins indicates no bias in sampling between untreated and LY-treated populations. **l.** Percent self-renewing^12^ cells with a cluster stability score ≥ 0.8 annotated as substates q3, q4, and q5 within the no BrdU condition across 50µm^2^ bins of increasing apical size.

By scRNAseq, *draxin* was enriched among cells highly responsive to short-term Notch perturbation, being q3 in subcluster identity^17,20^. When expression levels for all 8 transcripts detected by smFISH were visualized across the population (**Figure 6c**), *draxin* transcripts were graded and coincident with increasing *ascl1a* and *dla* expression, both associated with NSC activation^45,46^, and with elevated *pcna* levels. Together, these patterns of gene expression suggested a transcriptionally encoded continuum of graded quiescence depth within the self-renewing^8^ cluster. On this basis, we assigned molecular quiescence scores derived from principal component analysis (PCA) as a measure of relative quiescence depth (**Figures 6d and S7b**). PC1 values were scaled between 0-1, with scores equivalent to 1 indicative of deep quiescence, represented by low levels of *draxin*, *dla*, *ascl1a* and *pcna* expression, and 0 being shallow (**Figures 6d and 6e**). Apical cell size was graded along the same axis within self-renewing^8^ continuum, inversely related to molecular quiescence score (**Figures 6f and 6g**) and in agreement with previous observations from live-imaging linking large apical size and NSC activation propensity^1,2^.

To experimentally test whether large cells, low in molecular quiescence score, are indeed in a functionally shallow state of quiescence, NSPCs were treated in vivo for 24 hours with the gamma-secretase inhibitor LY411575 (LY) to perturb Notch signaling. We reasoned that if quiescence depth is inversely related to apical cell size, large cells within the self-renewing^8^ pool should be most sensitive to short-term Notch inhibition (**Figure 6h**). Following the treatment, gene expression was assessed in situ for the same 8 transcripts as before. Decreased *mfge8a* expression with elevated *ascl1a* and *dla* confirmed the efficacy of the treatment relative to a DMSO control (**Figures S7c and S7d**). 98 LY-treated cells were added to the integrated smFISH dataset (**Figure S11e**), with Gs intensities and apical cell sizes being comparable between untreated and LY-treated populations (**Figures S7f and S7g**), indicating no cell-type bias between conditions. We then assessed quiescence depth for self-renewing^8^ cells with cluster stability scores ≥ 0.8 (2522 cells total) (**Figure 6i**). The normal distribution of molecular quiescence depth among untreated cells was shifted to a bimodal distribution upon LY treatment due to an increase in the proportion of cells with quiescence scores ≤ 0.5 (**Figure 6j**). When cells were binned by apical area, quiescence scores were found to be higher among smaller cells (50-100 and 100-150µm^2^ bins), independent of treatment (**Figure 6k**). In contrast, Notch inhibition led to a dramatic decrease in quiescence score among large cells (150-200 and 200-250µm^2^ bins) compared to untreated cells, consistent with larger cells being shallower in their quiescence depth.

Finally, to compare molecular quiescence scores against scRNAseq subcluster identities, we returned to the “no BrdU” dataset alone for which scRNAseq q-subcluster annotations were successfully transferred (**Figure 2c**). Among self-renewing^12^ cells with cluster stability scores ≥ 0.8, we determined the percentage of q3, q4, and q5 annotations across apical size bins of 50µm^2^ increments. We found that the proportion of cells assigned to q3 (shallow quiescence) increased with apical size, particularly among cells greater than 100µm^2^ (**Figure 6l**).

Together, these data demonstrate that a transcriptional continuum between deep and shallow quiescence, independent of lineage progression, is contained within the self-renewing^8^ pool. Further, they also validate the use of a molecular quiescence score as a quantitative measure of relative quiescence depth, and underscore the link between transcriptional quiescence depth and apical cell size.

### Individual self-renewing NSCs transition through a transcriptionally encoded quiescence cycle oriented from deep to shallow states

Differences in quiescence depth may result from distinct NSC subpopulations that vary in proliferation frequency, thus differentially represented among labelled BrdU^pos^ progeny, or from dynamic trajectories by individual cells transitioning between deep and shallow states (**Figure 7a**). To distinguish between these possibilities, we exploited our BrdU time-stamping data. We focused on BrdU^pos^ self-renewing^8^ cells post-division. When examined relative to the self-renewing^8^ population as a whole, all BrdU-labelled cells were localized to one side of the transcriptional continuum (**Figure 7b**). The quiescence score for each of these cells was between 0.5-1, with no obvious temporal ordering across time points (**Figures 7c-e**), suggesting that NSCs return to deep states of quiescence post-division that are maintained for up to 28 days.

**Figure 7.**
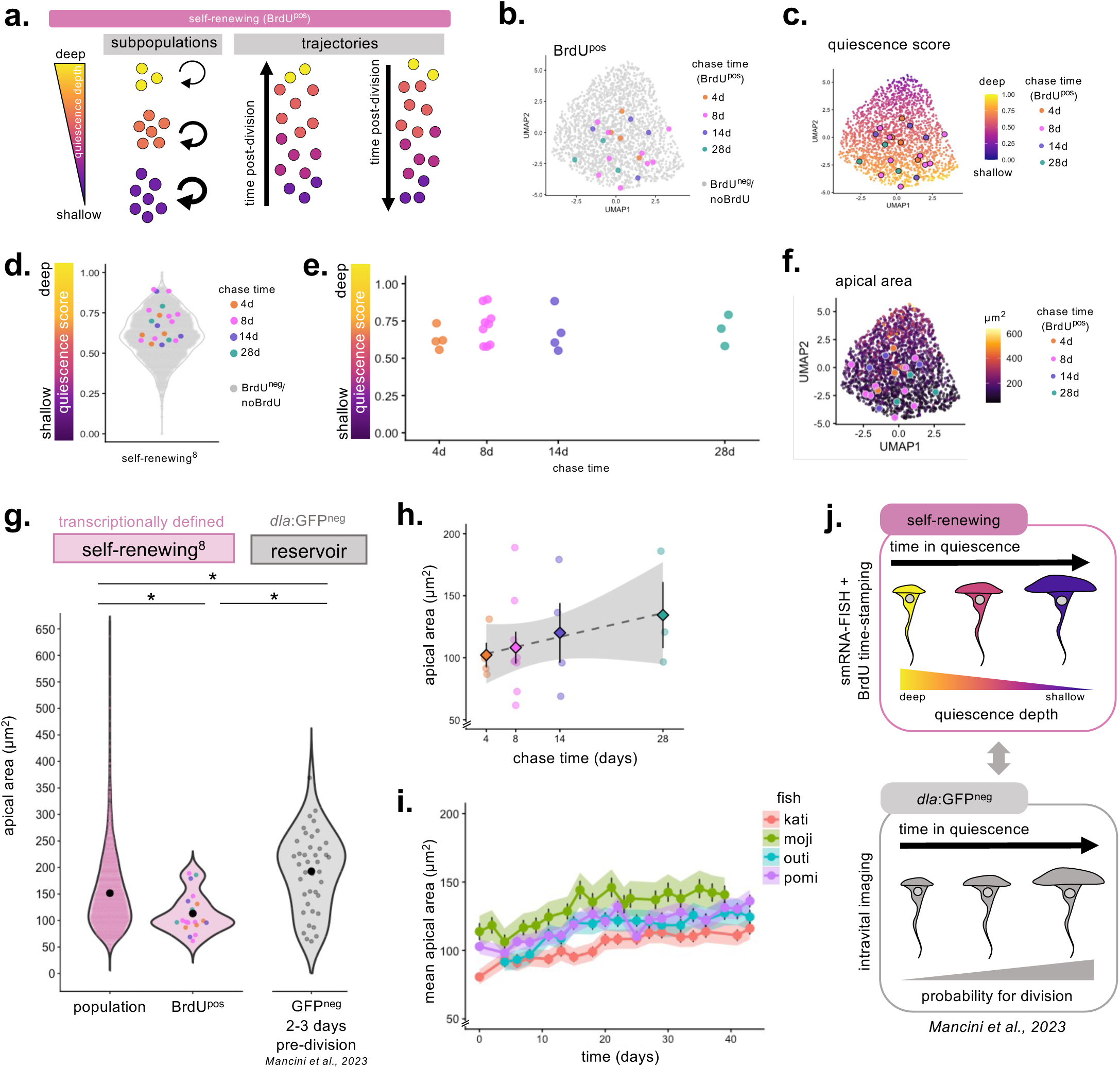
Self-renewing NSCs experience a molecular trajectory from deep to shallow quiescence coincident with a progressive increase in apical cell size. **a**. Transcriptional heterogeneity among self-renewing^8^ NSCs associated with differences in quiescence depth may result from subpopulations varying in their proliferation frequencies (left), indicated by the thickness of the circular arrows, or from transcriptional trajectories within quiescence by which cells transit from shallow to deep, or from deep to shallow, over time. **b.** BrdU^pos^ NSCs positioned within the self-renewing^8^ UMAP color-coded by chase time**. c, d.** Quiescence scores among 2055 self-renewing^8^ cells represented by UMAP (c) and by violin plot (d). BrdU^pos^ cells are overlayed and color-coded by chase time. **e.** Quiescence scores for BrdU-labelled self-renewing^8^ cells across increasing quiescence durations. **f.** Apical area values among 2055 self-renewing^8^ cells represented by UMAP. BrdU^pos^ cells are overlayed and color-coded by chase time. **g.** Apical cell size distributions across the population of 2055 transcriptionally defined self-renewing^8^ NSCs compared to 20 BrdU^pos^ self-renewing^8^ NSCs post-division (all chase times), and 40 GFP^neg^ mother cells (MCs) pre-division from four Tg(*dla*:GFP) pallia^1^. Mean apical area values for each population are indicated in black. Asterisks indicate statistically significant differences determined by Welch’s one-way ANOVA followed by Games-Howell post-hoc tests with Bonferonni correction adjusted for three comparisons. **h.** Apical cell size measurements for self-renewing^8^ BrdU^pos^ cells at increasing chase times post-division. Dashed line shows the best-fit linear regression (slope = 1.4µm^2^ per day with p = 0.2 by two-tailed t-test). Shaded region indicates the 95% confidence interval. Circles represent cells measured in situ. Diamonds represent the mean ± s.d. for each chase time. **i.** Mean apical area over time among non-dividing GFP^neg^ cells across four Tg(*dla:GFP*) adult pallia tracked for 43 days by live imaging. Shaded region represents the 95% confidence interval. Black vertical lines represent the standard error. **j.** A transcriptional trajectory within self-renewing^8^ NSCs by which cells transition from deep to shallow quiescence over time, coincident with an increase in apical cell size, in consistent with previous observations by live imaging that showed apical cell size increases over quiescence durations among dla:GFP^neg^ NSCs, with large cells having a greater probability of division^1^.

Among self-renewing NSCs, quiescence phases are estimated to be multiple months in their duration, extending well beyond the 28-day chase time considered here^1,2^. To explore whether individual self-renewing NSCs gradually progress toward shallow substates of quiescence during these prolonged phases, we first leveraged the relationship between quiescence depth and apical cell size. We compared the mean apical size among deeply quiescent BrdU^pos^ cells post-division to that of the entire self-renewing^8^ population (**Figures 7f and 7g**). As expected, BrdU-labelled cells were relatively small, having a mean apical area of 113 ± 8 µm^2^ compared to 152 ± 2 µm^2^ measured among all self-renewing^8^ cells (**Figure 7g**). Using previously recorded movies from Tg(*dla:GFP*) pallia in which cells were tracked by live imaging 2-3 days prior to division^1^, the mean apical area of GFP^neg^ (reservoir) mother cells (MCs) was also compared (**Figure 7g**), with such cells being similarly self-renewing and close to division. GFP^neg^ MCs were 193 ± 12 µm^2^, approximately twice the apical size of BrdU^pos^ self-renewing^8^ cells at 4-days post-division (102 ± 10 µm^2^), and considerably larger than the self-renewing^8^ population. Together, these values suggest that in order for daughter cells (post-division) to become MCs (pre-division), they must double their apical size during quiescence. Further, since apical size correlates with quiescence depth, and the mean apical size of MCs is considerably larger than that of the self-renewing^8^ population, only a minority of NSCs are likely to be in shallow quiescence at any given time.

Finally, we examined apical surface sizes among BrdU-labelled self-renewing^8^ cells at increasing chase times post-division, with such cells being NSCs experiencing increasing quiescence durations (**Figure 7h**). Values trended upward, albeit lacking statistical significance due to the high variability between cells at each chase time. For comparison, we similarly assessed the temporal progression of mean apical size among non-dividing GFP^neg^ cells over 43 days, previously measured by live imaging within four Tg(*dla:GFP*) pallia^1^. A progressive increase in apical cell size was observed over time, similar in magnitude to static measures using BrdU time-stamping (**Figure 7i**, compare with **7h**). Taken together, these data suggest that self-renewing NSCs post-division transit through the entire transcriptomic space as they progress from deep to shallow quiescence, tractable as a progressive increase in apical size (**Figure 7j**). This results in a quiescence cycle oriented from deep to shallow quiescence, leading to cell cycle entry, by which self-renewal at division is achieved.

## DISCUSSION

This work proposes an experimental framework for deciphering transcriptional trajectories of individual NSCs during state and fate decisions. Using BrdU time-stamping and a molecular code tractable in situ, we resolve cell state transitions among individual NSPCs. Such transitions pertain to division, lineage progression, and for self-renewing NSCs, molecular state changes during quiescence. Regarding lineage progression, we reveal that NSPCs are ordered along the main axis of the transcriptional continuum by stemness potential, separated into two differentially fated clusters, and that cells transition from the self-renewing to the committed cluster via cryptic division events. Among quiescent self-renewing NSCs, where transcriptional heterogeneities reflect variable quiescence depths, we demonstrate that individual NSCs experience a temporally ordered series of transcriptional substates. These substates form a quiescence cycle through which NSCs transit between divisions. Through this cycle, NSCs progressively explore the entire transcriptional space spanning from deep to shallow substates. These results resolve the main molecular trajectories of individual NSCs, revealing state and fate transitions that ensure stemness maintenance and continued neurogenesis in the adult vertebrate pallium. While regulation of molecular cell trajectories likely also occurs at post-transcriptional levels^47^, not analyzed here, our results show that transcriptional identities among individual NSCs capture key NSC decisions, establishing transcriptional identity as a primary readout of NSC state and fate.

### Transcriptional identity combined with real measures of time in situ are essential for resolving single-cell trajectories

Various computational approaches have been developed to resolve single-cell trajectories, relying on transcriptional proximities and rapid temporal dynamics to identify probable transitions over inferred pseudotime at population scale^22–26^. These approaches, however, produce trajectories that are difficult to interpret when applied to progressive cell-state transitions over extended timeframes, such as those among somatic adult stem cells, in which dynamic population homeostasis prohibits cross-time comparisons. To overcome this, we examined NSPC trajectories in situ, allowing for transcriptional identity to be assessed in the context of cell morphology, kin relationships, and real measures of time. Several methodological developments were required for this work. First, multiplexed smFISH needed to be adapted for whole-mount tissue, targeting a minimal subset of differentially expressed transcripts. Image analysis tools, like *Multireg* and *FishFeats*, were also required, designed to facilitate quantification of multiplexed smFISH data captured across multiple rounds of imaging^32^. Finally, a method for multimodal transcriptomic data integration was applied to transition between scRNAseq and smFISH modalities, a major bottleneck in current single-cell analyses. Combined, these tools enabled us to map scRNA-seq identities q1-q5 onto cells for which gene expression was measured in situ, allowing for the biological significance of transcriptional heterogeneities to be determined both at the population level and within individual lineages. Then, combined with BrdU-based time-stamping, these methods allowed for cell state transitions hidden within transcriptional heterogeneity to be uncovered, revealing two major trajectory-types: those that underly lineage progression punctuated by amplifying divisions, and those by which self-renewing NSCs transition between substates of variable quiescence depth.

### NSPC trajectories associated with lineage progression involve cell divisions hidden within the transcriptomic space

By coupling smFISH to BrdU-labelling, sister cells were tracked within transcriptional space. This demonstrated that the transcriptional continuum of quiescent pallial NSPCs is interrupted by a series of cell divisions, which had not been predicted by pseudotime analyses^20^. Further, it allowed for transcriptional identities of sister cells to be compared, revealing that most divisions amplify the committed population, being symmetric in fate. Rarer cases of fate asymmetry at division were also observed, indicated by immediate transcriptional asymmetry by which sister cells were partitioned between the self-renewing and committed clusters. Such divisions, originating from self-renewing mother cells, further highlight that self-renewing mother and daughter cells belong to the same transcriptional subspace, suggesting *bona fide* transcriptional self-renewal.

### Among self-renewing NSCs, transcriptional heterogeneities are associated with physiological variability in quiescence depth

An important outcome of our multimodal approach, by which scRNAseq with smFISH, IHC, and BrdU tracing are combined, is the identification of transcriptional signatures restricted to self-renewing NSCs. We find this population, represented here by the self-renewing^n^ clusters and equivalent to scRNA-seq subclusters q3-q5, to contain a transcriptional continuum within itself, independent of that encoding lineage progression. We propose that this continuum reflects cells transitioning through substates of variable quiescence depths, with progression along the quiescence depth continuum quantifiable as a molecular quiescence score. This interpretation is supported by several converging arguments. First, genes associated with NSC activation are graded in expression along this progression. These include *pcna*, required for DNA replication and induced in late G1 of the cell cycle^48,49^, and zebrafish homologs of *Dll1* (*dla*) and *Ascl1* (*ascl1a*), expressed in activated NSCs^45,46^. With Notch2/3 signaling known to promote adult NSC quiescence, Dla ligand expression in shallow NSCs may act to cis-inhibit the Notch receptor, transiently downregulating signaling to promote activation^50–55^. The increased expression of *ascl1a* in shallow quiescence further supports this model, with murine *ascl1* known to be a target of Notch-mediated repression and required for NSC proliferation^45,56–58^.

Second, combining smFISH with apical area measurements revealed an inverse relationship between molecular quiescence score and apical cell size, indicating large NSCs are shallower in their quiescence depth (**Figure 6d and 6f**). This is consistent with previous live-imaging data linking increased apical size with greater probability for division^1^, and with recent work describing elevated mTORC1 activity among self-renewing NSCs large in apical size^59^. Together, these observations provide further evidence to support for our transcriptionally defined quiescence score as a relevant measure of quiescence depth. As a functional validation, we found that self-renewing NSCs large in apical size (150-250µm^2^), and therefore predicted to be transcriptionally closer to activation, were indeed those most responsive to Notch blockade. Such cells underwent a pronounced shift in quiescence score, indicative of a molecular transition towards activation, compared to smaller cells within the same cluster that exhibited no such change (50-150µm^2^) (**Figure 6k**). Finally, annotation transfer showed that increased apical area was associated with an increased representation of the q3 substate (**Figure 6l**), previously shown to be sensitive to short-term Notch blockade, thus representing shallow quiescence, in contrast to q4-q5 that remained resistant to Notch perturbation and were therefore interpreted to represent deep^17^.

### Self-renewing NSCs transit through an ordered and transcriptionally encoded quiescence cycle between divisions

We find that quiescence heterogeneity is temporally ordered, reflecting a molecular progression oriented from deep to shallow quiescence through which NSCs transition over time. By combining BrdU time-stamping with smFISH, we directly demonstrate that immediately after division, self-renewing daughter NSCs populate deep quiescence. In addition, we find that this state is maintained for at least 28 days (**Figures 7a-e**). Based on further converging arguments, we propose that cells then transition through a series of molecular states by which they reach shallow quiescence. Apical area measured among self-renewing NSCs post-division revealed a gradual increase in apical size over time (**Figure 7h**), and NSCs in shallow quiescence defined by smFISH, as well as MCs pre-division observed by dynamic imaging, were among those largest in apical area (**Figures 6d, 6f and 6g**). Given the relationship between quiescence depth and apical cell size, these observations support a model in which self-renewing NSCs progressively explore the entire self-renewing transcriptomic continuum, reaching shallow quiescence over a quiescence cycle that extends beyond the 28-day time course considered here.

Based on the mean apical sizes of BrdU-labelled cells across chase times post-division, apical area was estimated to increase at an average rate of 1.4µm^2^ per day. At this growth rate, 73 days are required for daughter cells to reach the apical size of MCs, assuming that MCs are twice the size of daughters (being 204 ± 20 µm^2^ based on the mean apical size of daughters 4d post-labelling). This estimation of average quiescence durations is shorter than those in previous studies, in which quiescence durations greater than 120 days were predicted^1,2^. These differences are likely due to a combination of factors, including pallial territories considered in each study, modalities used to define the self-renewing pool, and how cell divisions were tracked. Across all three studies, however, quiescence phases in self-renewing pallial NSCs last for months, establishing a multi-month timescale for the quiescence cycle.

Differences in quiescence depth reflect a readiness to engage in the cell cycle, often revealed by variable activation speeds in response to a stimulus^16,17,60–63^. A quiescence cycle was previously postulated in muscle stem cells (MuSCs) based on the appearance of a shallow quiescence state under injury conditions^64^. Our results extend this notion to physiological stem cell trajectories and show that this cycle is encoded by a transcriptional progression. This cycle may function as an intrinsic cell-autonomous clock, by which cell preparedness for the next division can be measured, but the mechanisms that control both the molecular sequence and the tempo of this cycle remain to be determined. As NSC activation is regulated by a combination of inputs that are both cell-autonomous and niche-derived^33,65–68^, each of these inputs may converge on a common regulatory mechanism that modulates the rate at which cells progress through the quiescence cycle.

### Deep quiescence dominates the quiescence cycle, with depth and duration inversely related

Notably, we find quiescence depth and duration to be inversely related among pallial NSCs, with quiescence depth decreasing over time. This is in contrast with other systems in which increasing quiescence durations are often associated with increased quiescence depth, requiring greater stimulation to resume proliferation^16,60,61,63,69^. Such studies, however, do not address single-cell state changes under physiological conditions. Instead, differences in quiescence depth are measured across populations in response to non-physiological quiescence-inducing signals, such as changes in cell density, adhesion, and nutrient availability, or in pathological contexts like chronological ageing. With transcriptional signatures of quiescence known to be dependent on the initiating stimulus^60^, it may be that the inverse relationship between quiescence depth and duration described here is a feature of physiologically induced quiescence.

The transition from deep to shallow quiescence is progressive and requires a considerable gene panel to distinguish between substates, precluding the use of live imaging to measure the relative durations of deep and shallow phases. The relative durations of each phase, however, can be inferred by the proportion of cells observed in each substate, with longer phases expected to contain a greater proportion of the population at any given time. By annotation transfer, we find that 41% of self-renewing (q3-q5) cells were q3 in identity, closely matching the 40% of cells annotated to q3 by direct scRNAseq sub-clustering. In contrast, only 22% of self-renewing cells had low molecular quiescence scores (between 0-0.5), suggesting that these scores likely represent a subpopulation within q3 being most transcriptionally proximal to activation. Nevertheless, both metrics identified a larger representation of cells in deep quiescence compared to shallow. Assuming that all cells undergo a progressive transition towards activation, these observations suggest that shallow quiescence is a rather transient phase, with deep quiescence being the predominant state within the quiescence cycle.

## LIMITATIONS OF THE STUDY

In this study, males and females were not considered separately, preventing sex-related differences in the quiescence cycle from being addressed. In addition, measures of time were incorporated by BrdU-labelling, biasing our understanding of NSPC trajectories to those among dividing cells. If a subset of NSCs within the self-renewing pool exhibits an especially low division rate, it will not be represented here, nor will cells that do not transition through the quiescence cycle, should such cells exist. In addition, due to technical limitations in orienting whole-mount pallia during imaging, smFISH experiments are largely restricted to Dm. As such, potential regional specificity in pallial NSPC identities is not addressed. While region-specific signatures within pallial NSPCs were not previously identified in scRNAseq studies by our group^20,59^, such signatures have been described by others^14,70^ and thus warrant consideration in future studies. With Notch activity is known to be spatially distributed across the pallial monolayer^1,6,71^, and likely to underly transitions between quiescence substates, spatial patterns of variable quiescence depths are likely to be similarly distributed across the niche as NSCs transit through the quiescence cycle in an asynchronous yet coordinated manner. While not explored here, the tools developed for this work will allow for further insight on spatiotemporal coordination of NSPCs substates at the molecular level.

## RESOURCE AVAILABILITY

### Lead contact

Requests for further information and resources should be directed to and will be fulfilled by the lead contact, Laure Bally-Cuif.

### Materials availability

This study did not generate new unique reagents.

### Data and code availability

This paper analyzes existing, publicly available scRNAseq data, deposited in the GEO database under accession number GSE225863 and available at https://www.ncbi.nlm.nih.gov/geo/query/acc.cgi?acc=GSE225863. Single-cell smFISH gene expression data and custom codes for their analyses will be shared by the lead contact upon request. Any additional information required to reanalyze the data reported in this paper is available from the lead contact upon request.

## ACKNOWLEDGEMENTS

Thanks to Sebastien Bedu and Nathan Guibert for expert fish care, and to all members of the ZEN lab for insightful discussions over the course of the project. Thanks to the Department of Developmental and Stem Cell Biology at Institut Pasteur, and to Florian Muller and Christophe Zimmer for their expert advice and constructive exchanges. Thanks to Ludovic Arnold and Melissa Turan from BioTechne for their technical support. This work was funded by the ANR (ANR PRC QDynamics to LBC and Labex Revive ANR-10-LABX-0073), the Fondation pour la Recherche Médicale (EQU202203014636 to LBC), CNRS, INSERM, Institut Pasteur and the European Research Council (ERC) (AdG SyG 595 101071786 – PEPS to LBC). It was also supported by a government grant managed by the ANR under the France 2030 program, with reference numbers ANR-24-EXCI-0001, ANR-24-EXCI-0002, ANR-24-EXCI-0003, ANR-24-EXCI-0004, ANR-24-EXCI-0005. TF was a recipient of a Roux-Cantarini fellowship from Institut Pasteur. The work of JS was funded by the Inception program (Investissement d’Avenir grant ANR-16-CONV-0005 to LC), while the work of LC is funded by European Union (ERC StG, MULTIview-CELL, 101115618 to LC).

## AUTHOR CONTRIBUTIONS

Conceptualization, L.B.C., D.M., T.F.; data curation, T.F., J.S., N.D., I.F., D.M.; formal analysis, T.F., D.M. J.S., M.D., N.D.; funding acquisition, L.B.C.; investigation, T.F. I.F., J.S., D.M., M.D.; methodology, T.F., I.F., J.S., L.C., N.D., G.L., D.M.; project administration, L.B.C.; resources, L.B.C., N.D., L.C.; visualization, T.F.; validation, L.B.C., T.F., J.S., L.C.; supervision, L.B.C, T.F.; software, N.D., G.L.; writing – original draft, T.F., J.S.; writing – review and editing, L.B.C., L.C., and T.F.

## SUPPLEMENTAL FIGURE TITLES AND LEGENDS

**Supplemental Figure S1. Workflow for multiplexed smRNA-FISH combined with fluorescence immunohistochemistry in whole mount tissue and associated image analysis tools. a.** Workflow for multiplexed smRNA-FISH in whole mount pallia. 12 RNAscope Hiplex v2 probes were hybridized to mRNA targets and revealed in groups of three via four sequential rounds of imaging, with fluorophores associated to probes cleaved between imaging rounds. Fluorescence immunostaining for Sox2 and Zo1 was present in all rounds. Immunostaining against endogenous Gs was captured in a fifth round of imaging. b. *Multireg* allowed for multiple confocal whole mount images to be aligned. A, Anterior; P, Posterior. Scale bars, 100μm. **c.** With *fishfeats,* apical cell contours and mRNA puncta were segmented and quantified from aligned images. Box indicates magnified region shown in (d). A, anterior; P, posterior. Scale bar, 100µm. **d.** Magnification of Sox2 (nuclear) and Zo1 (apical cell contour) immunostaining with segmentation performed using the Zo1 signal (panel 1). mRNA puncta were segmented in 3D (panel 2) and assigned to corresponding cell surfaces by 2D projection (panel 3). Scale bars, 50μm. **e.** Schematic representation of mRNA assignment to cellular surfaces by 2D projection. Manual correction was required for cells with smaller apical surfaces and larger cell bodies nested below apical surfaces of neighbouring cells. Apical Zo1 contours shown in dark blue; segmented surfaces indicated in red, purple, and light blue; mRNA puncta represented by solid circles within cell bodies; 2D projected puncta are shown by dashed circles above. Arrows indicate mismatch between the outline of the projected mRNA and the color within for which manual correction was applied. **f.** Correlation matrix summarizing the relationship between transcripts measured *in situ*.

**Supplemental Figure S2. Gene expression and apical cell size measured in situ distinguish self-renewing NSCs from committed progenitors. a.** Dendogram from hierarchical clustering for 821 cells within one pallial hemisphere using gene expression values detected using the 12-gene transcriptomic code. **b.** Stability scores following hierarchical clustering visualized by UMAP. **c.** Representation of Gs intensities across clusters 1^12^ and 2^12^ expressed as percentages. Cluster 1^12^: 98.5% Gs^high^, 1.1% Gs^low^, 0.4% Gs^neg^; cluster 2^12^: 35.5% Gs^high^, 17.2% Gs^low^, and 47.3% Gs^neg^. **d.** Apical sizes within clusters 1^12^ and 2^12^. Asterisks indicate highly significant difference determined by Wilcoxon rank-sum test (p < 2.2 x 10^-16^). **e.** 3D reconstruction of a confocal image of a Tg*(dla:GFP*) whole mount pallium following immunostaining for Zo1, Sox2, Gs, and GFP. Variable Gs and GFP levels are indicated by arrowheads. Scale bars, 10µm. **f, g.** Distribution (f) and percentage (g) of Gs intensities by GFP category across two Tg(*dla:GFP*) pallia in which 404 and 304 cells were considered. Values in (g) represent mean ± s.d. **h.** 3D reconstruction of a confocal image following immunostaining of a Tg*(dla:GFP*) whole mount pallium for Zo1, Sox2, and GFP combined with smRNA-FISH for *mfge8a*. Representative GFP intensities are indicated by arrowheads. Zo1 contours of representative cells for each GFP category are outlined by dashed lines. Scale bars, 10µm. **i.** Distribution of apical area values measured among GFP^neg,^ GFP^weak,^ GFP^med^, and GFP^high^ cells across two Tg(*dla:GFP*) pallia (rep1 and rep2, minimum 500 cells per replicate). j. *mfge8a* expression within cluster 1^12^ and cluster 2^12^ from 821 cells measured by smRNA-FISH using the 12-gene transcriptomic code. Values indicate the number of transcripts per cell. Asterisks indicates highly significant difference (p < 2.2 x 10^-16^) determined using Wilcoxon rank-sum test. k. *mfge8a* expression measured among GFP^neg,^ GFP^weak,^ GFP^med^, and GFP^high^ cells across two Tg(*dla:GFP*) pallia (rep1 and rep2, minimum 500 cells per replicate). Values indicate the number of transcripts per cell.

**Supplemental Figure S3. Transcriptional asymmetry between BrdU-labelled sister cells within self-renewing^8^ cell-containing clones is detected 24h post-division.** a. *mfge8a*, *cspg5b*, *timp4.3*, *draxin*, *pcna*, Gs, and apical area among 435 NSPCs within a pallium 24h post-BrdU-labelling (10h). **b.** Hierarchical clustering of 435 NSPCs based on the expression of five genes measured by multiplexed smRNA-FISH identifies two major clusters, cluster 1^5^ (self-renewing) and cluster (committed) 2^5^. **c.** Cluster stability scores for individual cells represented by UMAP. **d.** Stability scores for BrdU-labelled sister cells among self-renewing cell-containing clones.

**Supplemental Figure S4. 8-gene smRNA-FISH datasets distinguishes self-renewing from committed cells within the integrated in situ dataset. a.** Pairwise comparison between *mfge8a*, *cxcl14*, and *slc4a4a*, and *timp4.3* and *igfbp2a* 821 cells measured in one hemisphere by smRNA-FISH. Values represent the number of transcripts per cell. **b.** Predicted accuracy of substate assignment using the combined expression of 8 genes (listed above) estimated by area under receiver-operator curves (AUC). **c.** Patterns of gene expression within the integrated dataset as visualized by UMAP. **d.** Dendogram following hierarchical applied to the integrated smRNA-FISH dataset. Boxes indicate self-renewing^8^ and committed^8^ clusters. **e.** Cluster stability scores for individual cells visualized by UMAP. **f.** Percentages of Gs^high^, Gs^low^, and Gs^neg^ cells within self-renewing^8^ and committed^8^ clusters. Cluster 1^8^: 93% Gs^high^, 6% Gs^low^, and 1% Gs^neg^; cluster 2^8^: 26% Gs^high^, 26% Gs^low^, and 48% Gs^neg^. **g.** Apical area distributions within self-renewing^8^ and committed^8^ clusters. Cluster 1^8^: 142 ± 69 µm^2^; cluster 2^8^: 40 ± 37µm^2^ (mean ± s.e.m.). h. *mfge8a* distributions across self-renewing^8^ and committed^8^ clusters. Cluster1^8^: 56 ± 26 transcripts per cell; cluster 2^8^: 10 ± 14 transcripts per cell (mean ± s.e.m.).

**Supplemental Figure S5. Fate asymmetry is observed among BrdU-labelled progeny within individual lineages at increasing chase times post-division. a, c, e, f.** 3D reconstructions of confocal images of self-renewing^8^ cell-containing BrdU^pos^ clones 4d (a), 8d (c), 14d (e), and 28d (f) post-labelling. White dashed lines indicate Zo1 (left) and BrdU (right) immunosignal for cells within BrdU^pos^ clones maintaining an apical surface revealed alongside Gs. At 8d post-division, Sox2 immunostaining was also performed. Grey dashed lines indicate BrdU^pos^ nuclei belonging to delaminated cells within the clone lacking an apical surface (not quantified). Arrowheads point to BrdU^pos^ nuclei from other clones. Delaminated BrdU^pos^ nuclei were assessed in optical cross section at 4d and 8d post-division (below), with white dashed lines indicating the nuclear contour of the self-renewing^8^ cell within the clone (Cell ID indicated above). Scale bars, 5µm**. b, d.** Lineage trees logically inferred by the number of BrdU^pos^ cells per clone, their apical sizes, and the relative intensity of BrdU-staining among cells within clones 4d (b) and 8d (d) post-division. Cell ID and apical cell sizes are indicated for all cells within the clone maintaining an apical surface.

**Supplemental Figure S6. Fate asymmetry among progeny within BrdU-labelled clones is visualized within transcriptomic space. a-h.** BrdU^pos^ cells within self-renewing^8^ cell-containing clones (a, c, e, g) and committed^8^ cell-only clones (b, d, f, h) at 4d (a, b), 8d (c, d), 14d (e, f), and 28d (g, h) post-labelling positioned within the integrated smFISH UMAP.

**Supplemental Figure S7 Molecular quiescence scores reflect a functional measure of quiescence depth in vivo. a.** Percentages of Gs^high^, Gs^low^, and Gs^neg^ cells among 2055 self-renewing^8^ cells with a cluster stability score ≥ 0.95. **b.** Gene loadings for PC1 and PC2 following a PCA using normalized *cspg5b*, *mfge8a*, *timp4.3*, *draxin*, *gyg1b*, *dla*, *ascl1a*, and *pcna* expression measured by multiplexed smRNA-FISH. **c.** Pallia were treated with either DMSO (control) or LY for 24h before *cspg5b*, *mfge8a*, *timp4.3*, *draxin*, *gyg1b*, *dla*, *ascl1a*, and *pcna* expression were measured by smRNA-FISH (left). A 3D reconstruction of a confocal image of these whole mount pallia shows smRNA-FISH combined with immunolabelling of Zo1 and Sox2 (right). Scale bar, 5µm. **d.** Distribution of *mfge8a*, *dla*, and *ascl1a* expression in DMSO- and LY-treated pallia expressed as the number of transcripts per cell. **e.** 98 LY-treated cells were integrated with untreated cells from the BrdU pulse-chase and no BrdU experiments (5016 cells) using normalized expression for *cspg5b*, *mfge8a*, *timp4.3*, *draxin*, *gyg1b*, *dla*, *ascl1a*, *pcna* was measured by smRNA-FISH and visualized by UMAP. **f.** Percentages of Gs^high^, Gs^low^, and Gs^neg^ cells among untreated and LY-treated pallia. **g.** Distribution of apical area among untreated and LY-treated cells showing no bias in cell size representation between datasets.

## KEY RESOURCES TABLE

| REAGENT OR RESOURCE | SOURCE | IDENTIFIER |
| --- | --- | --- |
| **Antibodies** |  |  |
| Monoclonal mouse-IgG1-anti-Zo1 | Fisher Scientific | 10129012 |
| Monoclonal mouse-IgG1-anti-Sox2 | Abcam | ab171380, RRID: AB_2732072 |
| Monoclonal mouse-IgG2a-anti-Gs | Sigma Aldrich | MAB302 |
| Polyclonal chicken-IgY-anti-GFP | Aves Labs | GFP-1020, RRID: AB_10000240 |
| Polyclonal rabbit-IgG-anti-BrdU | Abcam | ab152095, RRID: AB_2813902 |
| Goat-anti-mouse-IgG1-405 | BioLegend | 409109, RRID: AB_10642830 |
| Goat-anti-mouse-IgG2a-488 | Invitrogen | A21131 |
| Goat-anti-mouse-IgG2a-633 | Invitrogen | A21136 |
| Goat-anti-chicken-488 | Invitrogen | A11039 |
| Goat-anti-rabbit-405 | Invitrogen | A31556 |
| Goat-anti-rabbit-546 | Invitrogen | A11010 |
| **Chemicals, peptides, and recombinant proteins** |  |  |
| BrdU | Sigma Aldrich | B5002-100MG |
| Hydrogen peroxide (H<sub>2</sub>O<sub>2</sub>) | Sigma Aldrich | 216763-100ML |
| Dimethyl sulfoxide (DMSO) | Sigma Aldrich | D8418-100ML |
| LY411575 | Sigma Aldrich | SML0506-25MG |
| Methanol (MeOH) | Sigma Aldrich | 34860-2.5L-R |
| 10X phosphate buffered saline (PBS) | Fisher Scientific | 10649743 |
| 16% paraformaldehyde (PFA) | Fisher Scientific | 15828264 |
| Tween-20 | Sigma Aldrich | P9416-100ML |
| Formamide | Sigma Aldrich | 47671-250ML-F |
| 20X saline-sodium citrate (SSC) | Fisher Scientific | 15557044 |
| Hydrochloric acid (HCl) | Sigma Aldrich | 258148-100ML |
| Triton-X100 | Sigma Aldrich | X100-100ML |
| Normal goat serum | Sigma Aldrich | G6767-100ml |
| **Critical commercial assays** |  |  |
| RNAscope™ HiPlex v2 assay | Bio-Techne Sales Corp | 324410 |
| 1X RNAscope™ Target Retrieval Reagent | Bio-Techne Sales Corp | 322000 |
| **Deposited data** |  |  |
| scRNAseq data | Morizet et al., 2024; GEO database | GEO accession GSE225863 |
| smFISH data | This paper | *Will be added to a repository upon acceptance* |
| **Experimental models: Organisms/strains** |  |  |
| *Danio rerio* AB/AB wild type strain | European Zebrafish Resource Center (EZRC) | https://zfin.org/ZDB-GENO-960809-7 |
| *Danio rerio* Tg(*dla:gfp*) | Madelaine and Blader, 2011 | https://doi.org/10.1016/j.ydbio.2010.10.025 |
| **Software and algorithms** |  |  |
| pROC (R) | Robin et al., 2011 | http://www.biomedcentral.com/1471-2105/12/77/ |
| sva (R) | Johnson et al., 2007;<br>Leek et al., 2012 | https://bioconductor.org/packages/sva |
| umap (R) | Konopka, 2023 | [10.32614/CRAN.package.umap](https://doi.org/10.32614/CRAN.package.umap) |
| stats (R) | R Core Team | https://www.R-project.org |
| ggplot2 (R) | Wickham, 2016 | https://ggplot2.tidyverse.org |
| ggcorrplot (R) | Kassambara, 2023 | https://doi.org/10.32614/CRAN.package.ggcorrplot |
| ggpubr (R) | Kassambara, 2026 | https://doi.org/10.32614/CRAN.package.ggpubr |
| dplyr (R) | Wickham, 2026 | https://doi.org/10.32614/CRAN.package.dplyr |
| emmeans (R) | Lenth, 2026 | https://CRAN.R-project.org/package=emmeans |
| SciPy1.0 (Python) | Virtanen et al., 2020 | https://doi.org/10.1038/s41592-019-0686-2 |
| scanpy v1.9.2 (Python) | Wolf, Angerer, and Theis, 2018 | https://doi.org/10.1186/s13059-017-1382-0 |
| ZEN blue v3.8 | ZEISS | https://www.microshop.zeiss.com/en/de/softwarefinder/software-categories/zen-black/ |
| Imaris v10 | Oxford Instruments | https://imaris.oxinst.com/ |
| Imaris stitcher | Oxford Instruments | https://imaris.oxinst.com/products/imaris-stitcher |
| Napari 0.4.19 | The napari team | https://napari.org/stable/index.html |
| Multireg | This paper | NA |
| FishFeats | Letort et al., 2025 | https://doi.org/10.1093/bioinformatics/btag105 |
| **Other** |  |  |
| Zeiss LSM980 upright confocal microscope | Zeiss | https://www.zeiss.com/microscopy/en/products/light-microscopes/confocal-microscopes.html |
| Polysine™ Adhesion microscope slides | Fisher Scientific | 10219280 |
| 22mm x 22mm #1 cover slip | Thorlabs | CG15CH2 |
| 0.5mm iSpacer | Nikon | 2SUN1016 |
| Immersol immersion oil | Zeiss | 433802-9010-000 |

## EXPERIMENTAL MODEL AND STUDY PARTICIPANT DETAILS

### Zebrafish

All procedures relating to zebrafish (*Danio rerio*) care and treatment conformed to the directive 2010/63/EU of the European Parliament and of the council of the European Union. Wild type AB/AB and Tg(*dla:gfp*) zebrafish^37^ were used for this study, aged 3-4 months post-fertilization without distinguishing between sexes. Fish were maintained in 3.5L tanks in 28.5°C and pH 7.4 water with a maximum density of 5 fish per L, as previously described^13^. Fish were euthanized in ice water for 10-minutes, according to the guidelines of the Ministry of Education, Research, and Innovation.

## METHOD DETAILS

### Transcriptomic code and subcluster prediction

Receiver-operator curves were used to estimate the sensitivity versus specificity of either the 8 or 12 gene transcriptional code, with area under the curve (AUC) reflecting confidence intervals calculated using the R package *pROC*^73^. Monte Carlo simulations were used to compare AUC values for 1000 receiver-operator simulations using random combinations of 12 genes among those detected by scRNAseq.

### BrdU labelling

To mark proliferating cells, zebrafish were placed in 1L beakers containing 1mM BrdU prepared as a 500µM stock in DMSO and diluted in 500ml fish water maintained at 28°C with continuous air flow for 10 hours. Following the treatment, fish were transferred to beakers containing 1L fish water for up to 48 hours, with one feeding and water change per day. Fish were then returned to 15L tanks and maintained in the animal facility for the remainder of the chase time before euthanasia and dissection. Chase times were 24h, 4 days, 8 days, 14 day, and 28 days.

### Pharmacological Notch inhibition

Transient inhibition of Notch activity was achieved by treating zebrafish with the γ-secretase inhibitor LY411575 prepared at a stock concentration of 100µM in DMSO and stored at -20°C shielded from light. LY411575 was diluted to a final concentration of 10µM in 500ml fish water. During treatment, fish were maintained at 28°C with continuous air flow for 24 hours, alongside controls treated with an equal volume of DMSO. Following treatment, fish were rinsed twice in fish water before euthanasia and dissection.

### Tissue isolation, fixation, and processing

Whole brains were dissected from adult zebrafish and fixed overnight at 4°C with rocking in 4% paraformaldehyde (PFA) prepared in 1 X PBS supplemented with 0.1% Tween-20 (PBS-T). Samples were then washed in PBS-T and dehydrated through a series of 5-minute PBS-T washes increasing in methanol concentration (25% / 50% / 75% / 100%) performed at room temperature, then stored at -20°C in 100% methanol. For processing, tissues were rehydrated using the reserves series of washes with methanol in PBS-T, followed by a final PBS-T wash. Samples were incubated in 1ml of pigment removal solution (3% H_2_O_2_, 5% formamide, and 0.5X SSC prepared in water) and incubated for 15-minutes under light. Brains were then washed in PBS-T for 5-minutes at room temperature with gentle rocking.

For BrdU-labelled samples, antigen retrieval was performed before rehydration, with samples first incubated in 0.2N HCl diluted in 100% methanol from a 2N stock for 30-minutes at room temperature with rocking. Samples were then rinsed with 100% methanol and rehydrated using a series of methanol washes, each being 5-minutes at room temperature with rocking and decreasing in concentration (75% / 50% / 25%), prepared in PBS-T. Following a final wash in PBS-T alone, tissues were incubated with 1X RNAscope^TM^ Target Retrieval Reagent, diluted to 1X in water from a 10X stock, at 98°C for 5-minutes with agitation.

### Single-molecule mRNA fluorescence *in situ* hybridization (smFISH)

To detect gene expression in whole mount pallia, the RNAscope^TM^ HiPlex v2 assay was performed according to the manufacturer’s instructions with some adaptations for whole mount tissue. All RNAscope^TM^ HiPlex v2 reagents were warmed to room temperature before use. Following rehydration and pigment removal, samples were placed in eppendorfs with 50μl of RNAscope^TM^ HiPlex v2 diluent buffer and pre-hybridized for 3 hours at 40°C with agitation. 1μl of each probe was diluted at 1:50 in diluent buffer to a final volume of 50μl (referred to hereafter as probe solution), then warmed to 40°C for 10 minutes. Diluent buffer was removed from tissue samples and replaced with probe solution, then incubated at 40°C overnight with agitation. Probe solution was then removed and samples were washed two times for 10 minutes in 1ml of 1X RNAscope^TM^ wash buffer at room temperature with gentle rocking, then incubated in 50μl of Amp1 for 30 minutes at 40°C with agitation, followed by two 10-minute washes in 1ml 1X RNAscope^TM^ wash buffer at room temperature. The same incubation and wash steps were repeated for Amp2 and Amp3. 1X RNAscope^TM^ wash buffer was then replaced with 50μl of FluoroT1-T3 and samples were incubated at 40°C for 15 minutes with agitation. Samples were washed three times in 1ml of 1X RNAscope^TM^ wash buffer for 10 minutes each time, followed by immunostaining (below), then mounted onto microscope slides for imaging (referred to as Round 1). Following image acquisition, tissue was removed from the slide and washed in PBS-T for 5 minutes at room temperature. A second 5-minute wash followed, this time with 4X SSC. Samples were then incubated for 15 minutes with gentle rocking at room temperature in 10% RNAscope^TM^ HiPlex V2 cleaving solution diluted in freshly prepared in 4X SSC. A 5-minute wash in PBS-T was performed before a second incubation in 10% cleaving solution, again for 15 minutes at room temperature with gentle rocking. Tissue was then transferred to a 15ml falcon tube and washed five times in 14ml of PBS-T with gentle rocking at room temperature. Samples were then transferred back to an Eppendorf and incubated with 50μl of FluroT4-T6 for 15 minutes at 40°C with agitation. Samples were washed three times in 1ml of 1X RNAscope^TM^ wash buffer for 10 minutes each time and mounted onto microscope slides for imaging (Round 2) with care to place the pallial hemisphere in a similar orientation as captured in the first image. Following image acquisition, the tissue was removed from the slide, transferred to an Eppendorf, and stored in PBS-T at 4°C overnight. The following morning, a 5-minute was in 4X SSC was performed, followed by the same cleaving and fluorophore hybridization steps as described for FluoroT4-T6 until probes associated with FluoroT7-T9 (Round 3) and FluoroT10-T12 (Round 4) were revealed and imaged. Quantifications for each transcript revealed using the 12-probe multiplexing protocol were compared to experiments in which probes were detected in a single round, in the absence of any cleaving, to confirm that gene expression counts in whole mount tissue were not impacted by fluorophore cleaving solution (data not shown).

### Immunostaining

For protein detection in situ, samples were incubated in 1ml of blocking solution (0.1% Triton X-100, 4% normal goat serum, and 0.1% DMSO prepared in 1X PBS) for 3 hours at room temperature with rocking. Primary antibodies were diluted in 500μl of blocking solution and incubated at 4°C overnight at the following dilutions: mouse-IgG1-anti-Zo1, 1:250; mouse-IgG1-anti-Sox2, 1:250; mouse-IgG2a-anti-Gs, 1:1000; chicken-anti-GFP, 1:1000; rabbit-anti-BrdU, 1:1000. The following day, samples were washed three times with 1ml of PBS-T at room temperature with rocking, then incubated in secondary antibodies diluted to 1:1000 in 500μl blocking solution at 4°C overnight. The following day samples were again washed three times with 1ml PBS-T before imaging. Due to their differential spatial localization, immunostaining for Zo1 and Sox2 were revealed in the same channel when combined with smFISH.

### Confocal microscopy

Whole mount images of telencephala were acquired using an upright Zeiss LSM980 Axio Examiner confocal microscope equipped with a 40X oil objective and ZEN black software. Samples were mounted within plastic spacers 0.5mm in thickness adhered to Polysine^TM^ Adhesion microscope slides and covered with a 22mm x 22mm #1 cover slip. A z-step of 0.4μm across a 15-20µm range, a resolution of 0.15-0.25um/pixel in xy, a numerical aperture of 1 airy unit (AU) for each channel with 2X averaging per line, 1024 × 1024-pixel resolution, and a bit-depth of 16 bits. Laser power was maintained between 0.8-2% for each experiment with detector gain adjusted according to the quality of staining.

### Image analysis

Image analysis was conducted in both Imaris (v10) and Napari (v0.4.19) using Python (v3.9).

#### Identification of BrdU-labelled nuclei

To assign BrdU^pos^ nuclei to the appropriate apical surface in the absence of additional nuclear markers, various cellular features were considered in 3D reconstructions of confocal image views in Imaris. These were: distance between BrdU^pos^ nuclei and apical cell surface, BrdU intensity, nuclear orientation, and continuity of BrdU immunosignal within the inferred nuclear volume deduced by the absence of cytosolic Gs immunostaining, carefully examined both in 3D reconstructions and 2D z-stacks for each BrdU^pos^ cell. Cells with discontinuous Zo1 staining, or for which not all 8 transcripts could be measured due to changes in tissue orientation between images, were not considered. Each cell was assigned an alpha-numeric Cell ID derived from the order in which the cell was segmented during image analysis preceded by the hemisphere in which the cell was observed (ie: RH16, cell 16 within the Right Hemisphere). Each BrdU^pos^ clone was also assigned a numerical Clone ID to track kin relationships.

#### Multiplexed smFISH image registration with Multireg

The napari plugin *multireg* was developed for the registration of 3D images of one field of view and with one color channel common to all images (in our case, Zo1 staining). It registers and corrects for changes in tissue shape and orientation that can occur as the tissue is unmounted from, and remounted onto, a microscope slide during repeated rounds of probe hybridization. For each image, the alignment and transformation are calculated with the python registration library itk-Elastix^40^ on the common channel that should be identical and then applied to all the other channels. First, an affine registration based on landmark points (manually placed on both the reference and moving images) was applied to globally align the images, bringing together similar regions. This first step allowed to identify the common region and correct for huge displacements/rotations between the mounted images. Then the registration was automatically refined with a B-spline based registration. This second step performed a finer registration also correcting for tissue deformation during the experimental procedure. Finally, the calculated transformation was applied to all the other color channels of the moving image. This procedure was repeated for the 3-4 stacks (unmounting/mounting iterations) to align. The final output of the plugin was one composite image containing the common channel, averaged once aligned over all or some of the iterations, and all other channels after registration. This allowed to then correctly quantify smFISH from multiplexed smFISH experiments in which different transcripts were sequentially imaged in the same region of whole mount tissue. *Multireg* is open-source, available on gitlab (gitlab.pasteur.fr/gletort/multireg) and can be installed as a pip module (pip install multireg) or within Napari in the Plugins interface.

#### Gene expression and cellular features in situ

The composite 14-channel image produced by *Multireg* was analyzed using a second napari plug-in, *FishFeats* ^32^. This custom plug-in proposes a flexible pipeline to perform analysis of 3D images of cells stained for junctions (Zo1), nuclei (DAPI) and smFISH. The junction and nuclei staining, acquired in the same channel, were separated by morphological filtering (top-hat based), a neural network specifically trained to discriminate between linear (junctions) and large volume (nuclei) structures (see https://gitlab.pasteur.fr/gletort/fishfeats/-/wikis/Separate-junctions-and-nuclei). The extracted junction staining was projected in 2D with a local max projection and segmented with EPySeg^74^. The segmentation was manually corrected, when necessary, through specific napari plug-in options to ease the manual correction. smFISH signal was segmented with Big-FISH^42^ using the automated threshold estimation. The segmented spots were then initially assigned to the cell which apical area was vertically above it (direct projection). Several initial assignment methods (see the options in the *FishFeats* plugin) were tested and the Z-projection gave the best initial assignment for most mRNAs. Spots that were too far from any apical area (too low or to the side) were considered unassigned. This initial assignment was then manually corrected through a specific option in the plugin, so that each segmented RNA spot was assigned to its most probable cell. Spots that were present in several color channels at once (called “overlapping spots” in *FishFeats*), likely to be staining artifacts, were unassigned. The number of smFISH dots per cell was quantified for each mRNA puncta, along with other cell features like apical cell area was also measured with the *FishFeats* plugin following segmentation. Cytoplasmic immunostaining intensity by manual assignment of cells to user defined categories (option “ClassifyCells” in *FishFeats*).

### Multimodal data integration by annotation transfer

scRNAseq data was preprocessed using the python package *scanpy v1.9.2*^75^. This preprocessing consists in three steps: first normalizing each cell by total counts over all genes, then log-transforming the data and lastly selecting highly variable genes using scanpy.pp.highly_variable genes with parameters: “min_mean=0.0125, max_mean=3, min_disp=0.5”. The Pearson correlation distance was then computed with the python package *scipy v1.10.1*^76^ between each scRNA and smFISH cell by using the 12 genes measured in both experiments. Using this distance matrix, the cluster annotations were transferred from the scRNA data to the smFISH cells using 20-nearest neighbor classification. In more detail, we associated to each smFISH cell a label corresponding to the most frequent cluster annotation among the 20 closest scRNA cells according to our distance matrix. 817 of 821 cells were successfully assigned q-subcluster identities, with the remaining 4 cells having exceptionally low levels of expression for each of the 12 genes assayed, and therefore corresponding to none of the possible q-subcluster annotations.

### Integration of BrdU-labelled smFISH datasets

For smFISH applied with 12 probes in the absence of BrdU (no BrdU), 821 cells across one pallial hemisphere were quantified. A second pallial hemisphere for which more than 200 cells were measured was similarly considered to ensure reproducibility (data not shown).

For smFISH applied following a 10-hour BrdU pulse and 24-hour chase, only one pallium was considered, with gene expression quantified among BrdU-labelled cells from both hemispheres and unlabelled cells from one hemisphere so as to generate a transcriptional continuum into which labelled cells could be positioned. 435 cells were considered in total.

For smFISH applied to pallia dissected from zebrafish pulsed for 10-hours with BrdU and followed by 4-, 8-, 14-, and 28-day chase times, two biological replicates, being pallia collected from two separate fish, were considered for BrdU-labelled samples 4 days (461 and 863 cells), 8 days (621 and 512 cells), and 28 days (626 and 431 cells) post-labelling, with only one pallium considered for the 14-day chase (775 cells). Gene expression values from each of these experiments, along with the no BrdU condition, were concatenated, log transformed, and scaled around 0 before batch correction was performed using the *ComBat* function in the *sva* R package^77,78^, selected after comparing between different batch correction methods. 2D UMAP embeddings for smFISH cell profiles using the *umap* R^79^ package with n_neighbors = 50 and min_dist = 0.5 to prioritize preservation of global data structure over local neighborhoods, using Euclidean distance metrics with a fixed random seed (123) to ensure reproducibility.

### Hierarchical clustering

Hierarchical clustering was performed on in situ gene expression in R using the base *stats* package using Euclidean distances and Ward’s method (Ward D2). For experiments in which either 12 or 5 probes were used to measure gene expression, clustering was performed on raw gene expression values. For 8-probe experiments in which multiple datasets were integrated with batch correction, log transformed and scaled values were used. Cluster stability was evaluated by bootstrapping the expression data 100 times with replacement. For each bootstrap sample, cells were hierarchically clustered and pairwise co-clustering frequencies recorded across iterations to produce a cell-by-cell stability matrix, normalized by the number of times each pair co-appeared in the resampled data. The resulting matrix represents the probability that any two cells are assigned to the same cluster under resampling. Per-cell stability scores were then calculated as the mean co-clustering frequency with other cells from the same original cluster.

### Quiescence scores

Quiescence scores were derived from principal component 1 (PC1) values following a principal component analysis (PCA) using the R base *stats* package. PC1 values were then scaled between 0 and 1 such that lower values represent cells in shallower quiescence depths.

### Data visualization

Barplots, violin plots, box-and-whisker plots, and dotplots were created with the R *ggplot2* package^80^ and the *ggpubr* extension^81^. Correlation matrices were created with the R package *ggcorrplot*^72^ on scaled and normalized gene expression. 2D UMAP embeddings generated for 12- and 8- probe smFISH experiments were obtained using the R package *umap*^79^ with n_neighbors = 75 and 50, respectively, with a min_dist = 0.5 applied to both experimental conditions, prioritizing preservation of global data structure over local neighborhoods. For experiments in which only 5 genes were detected, n_neighbors = 100 and min_dist = 0.9 were used. In all cases, Euclidean distance metrics were applied with a fixed random seed (123) to ensure reproducibility. Lineage trees were created with the R package *ape*^82^ for BrdU-labelled clones containing cells with apical domains for which all labelled nuclei (both within and below the NSPC layer) could be reliably identified. UMAP coordinates were then plotted with the R *ggplot2* package^80^.

To visualize the relationship between apical size and quiescence score across increasing apical size bins with and without LY treatment, analysis was restricted to cells with apical sizes between 50-250µm^2^, with 50µm^2^ being minimum size among self-renewing^8^ cells, and larger cells being outliers (sizes > 1.5X interquartile range). Within each size bin, kernel density estimates (KDE) were plotted for LY-treated and untreated cells. Curves were smoothed using a Gaussian kernel with a bandwidth adjustment factor of 1.2, with plots generated using the R packages *dplyr*^83^ and *ggplot2*^80^.

To estimate increases apical size over time, linear regressions were performed with the R base *stats* package.

## QUANTIFICATION AND STATISTICAL ANALYSIS

### Statistical analyses

Details regarding mean comparisons, error bars, and statistical tests can be found in the figure legends. Welch’s one-way ANOVA was performed followed by Games-Howell post-hoc tests with Bonferonni correction adjusted for three comparisons. Two-sided Wilcoxon rank-sum tests were performed with the base R *stats* package. Estimating least-squares means was performed across experimental conditions using the *emmeans* R package^84^, with pairwise differences between datasets assessed with Tukey-adjusted contrasts. For all statistical tests, significance was defined as p < 0.05.

## REFERENCES

1. Mancini, L., Guirao, B., Ortica, S., Labusch, M., Cheysson, F., Bonnet, V., Phan, M.S., Herbert, S., Mahou, P., Menant, E., et al. (2023). Apical size and deltaA expression predict adult neural stem cell decisions along lineage progression. Sci. Adv. 9. 10.1126/sciadv.adg7519.

2. Than-Trong, E., Kiani, B., Dray, N., Ortica, S., Simons, B., Rulands, S., Alunni, A., and Bally-Cuif, L. (2020). Than-Trong et al Lineage hierarchies and stochasticity ensure the long-term maintenance of adult neural stem cells.

3. Bottes, S., Jaeger, B.N., Pilz, G.A., Jörg, D.J., Cole, J.D., Kruse, M., Harris, L., Korobeynyk, V.I., Mallona, I., Helmchen, F., et al. (2021). Long-term self-renewing stem cells in the adult mouse hippocampus identified by intravital imaging. Nat. Neurosci. 24, 225–233. 10.1038/s41593-020-00759-4.

4. Ehm, O., Göritz, C., Covic, M., Schäffner, I., Schwarz, T.J., Karaca, E., Kempkes, B., Kremmer, E., Pfrieger, F.W., Espinosa, L., et al. (2010). RBPJκ-dependent signaling is essential for long-term maintenance of neural stem cells in the adult hippocampus. Journal of Neuroscience 30, 13794–13807. 10.1523/JNEUROSCI.1567-10.2010.

5. Barbosa, J.S., Sanchez-Gonzalez R, Di Giaimo R, Baumgart E V, Theis F J, Götz M, and Ninkovic J (2015). Live imaging of adult neural stem cell behavior in the intact and injured zebrafish brain. Science (1979). 348, 789–793. 10.1126/science.aaa2729.

6. Dray, N., Mancini, L., Binshtok, U., Cheysson, F., Supatto, W., Mahou, P., Bedu, S., Ortica, S., Than-Trong, E., Krecsmarik, M., et al. (2021). Dynamic spatiotemporal coordination of neural stem cell fate decisions occurs through local feedback in the adult vertebrate brain. Cell Stem Cell 28, 1457–1472.e12. 10.1016/j.stem.2021.03.014.

7. Rothenaigner, I., Krecsmarik, M., Hayes, J.A., Bahn, B., Lepier, A., Fortin, G., Götz, M., Jagasia, R., and Bally-Cuif, L. (2011). Clonal analysis by distinct viral vectors identifies bona fide neural stem cells in the adult zebrafish telencephalon and characterizes their division properties and fate. Development 138, 1459–1469. 10.1242/dev.058156.

8. Pilz, G.-A., Bottes, S., Betizeau, M., Jörg, D.J., Carta, S., April, S., Simons, B.D., Helmchen, F., and Jessberger, S. Live imaging of neurogenesis in the adult mouse hippocampus.

9. Dray, N., Bedu, S., Vuillemin, N., Alunni, A., Coolen, M., Krecsmarik, M., Supatto, W., Beaurepaire, E., and Bally-Cuif, L. (2015). Large-scale live imaging of adult neural stem cells in their endogenous niche. Development (Cambridge) 142, 3592–3600. 10.1242/dev.123018.

10. Cebrian-Silla, A., Nascimento, M.A., Redmond, S.A., Mansky, B., Wu, D., Obernier, K., Rodriguez, R.R., Gonzalez-Granero, S., Garcia-Verdugo, J.M., Lim, D.A., et al. (2021). Single-cell analysis of the ventricular-subventricular zone reveals signatures of dorsal and ventral adult neurogenic lineages. Elife 10. 10.7554/eLife.67436.

11. Cebrian-Silla, A., Nascimento, M.A., Mancia, W., Gonzalez-Granero, S., Romero-Rodriguez, R., Obernier, K., Steffen, D.M., Lim, D.A., Garcia-Verdugo, J.M., and Alvarez-Buylla, A. (2025). Neural stem cell relay from B1 to B2 cells in the adult mouse ventricular-subventricular zone. Cell Rep. 44. 10.1016/j.celrep.2025.115264.

12. Shin, J., Berg, D.A., Zhu, Y., Shin, J.Y., Song, J., Bonaguidi, M.A., Enikolopov, G., Nauen, D.W., Christian, K.M., Ming, G.L., et al. (2015). Single-Cell RNA-Seq with Waterfall Reveals Molecular Cascades underlying Adult Neurogenesis. Cell Stem Cell 17, 360– 372. 10.1016/j.stem.2015.07.013.

13. Mitic, N., Neuschulz, A., Spanjaard, B., Schneider, J., Fresmann, N., Novoselc, K.T., Strunk, T., Münster, L., Olivares-Chauvet, P., Ninkovic, J., et al. (2024). Dissecting the spatiotemporal diversity of adult neural stem cells. Mol. Syst. Biol. 20, 321–337. 10.1038/s44320-024-00022-z.

14. Cosacak, M.I., Bhattarai, P., Reinhardt, S., Petzold, A., Dahl, A., Zhang, Y., and Kizil, C. (2019). Single-Cell Transcriptomics Analyses of Neural Stem Cell Heterogeneity and Contextual Plasticity in a Zebrafish Brain Model of Amyloid Toxicity. Cell Rep. 27, 1307–1318.e3. 10.1016/j.celrep.2019.03.090.

15. Rigo, P., Ahmed-de-Prado, S., Johnston, R.L., Choudhury, C., Guillemot, F., and Harris, L. (2025). Sequential transcriptional programs underpin activation of hippocampal stem cells.

16. Harris, L., Rigo, P., Stiehl, T., Gaber, Z.B., Austin, S.H.L., Masdeu, M. del M., Edwards, A., Urbán, N., Marciniak-Czochra, A., and Guillemot, F. (2021). Coordinated changes in cellular behavior ensure the lifelong maintenance of the hippocampal stem cell population. Cell Stem Cell 28, 863–876.e6. 10.1016/j.stem.2021.01.003.

17. Morizet, D., Foucher, I., Mignerey, I., Alunni, A., and Bally-Cuif, L. (2025). Notch signaling blockade links transcriptome heterogeneity in quiescent neural stem cells with reactivation routes and potential.

18. Rigo, P., Ahmed-de-Prado, S., Johnston, R.L., Choudhury, C., Guillemot, F., and Harris, L. (2025). Sequential transcriptional programs underpin activation of hippocampal stem cells.

19. Dulken, B.W., Leeman, D.S., Boutet, S.C., Hebestreit, K., and Brunet, A. (2017). Single-Cell Transcriptomic Analysis Defines Heterogeneity and Transcriptional Dynamics in the Adult Neural Stem Cell Lineage. Cell Rep. 18, 777–790. 10.1016/j.celrep.2016.12.060.

20. Morizet, D., Foucher, I., Alunni, A., and Bally-Cuif, L. (2024). Reconstruction of macroglia and adult neurogenesis evolution through cross-species single-cell transcriptomic analyses. Nat. Commun. 15. 10.1038/s41467-024-47484-1.

21. Tritschler, S., Büttner, M., Fischer, D.S., Lange, M., Bergen, V., Lickert, H., and Theis, F.J. (2019). Concepts and limitations for learning developmental trajectories from single cell genomics. Preprint at Company of Biologists Ltd, 10.1242/dev.170506 https://doi.org/10.1242/dev.170506.

22. Lange, M., Bergen, V., Klein, M., Setty, M., Reuter, B., Bakhti, M., Lickert, H., Ansari, M., Schniering, J., Schiller, H.B., et al. (2022). CellRank for directed single-cell fate mapping. Nat. Methods 19, 159–170. 10.1038/s41592-021-01346-6.

23. Bergen, V., Lange, M., Peidli, S., Wolf, F.A., and Theis, F.J. (2020). Generalizing RNA velocity to transient cell states through dynamical modeling. Nat. Biotechnol. 38, 1408–1414. 10.1038/s41587-020-0591-3.

24. Qiu, X., Mao, Q., Tang, Y., Wang, L., Chawla, R., Pliner, H.A., and Trapnell, C. (2017). Reversed graph embedding resolves complex single-cell trajectories. Nat. Methods 14, 979–982. 10.1038/nmeth.4402.

25. Street, K., Risso, D., Fletcher, R.B., Das, D., Ngai, J., Yosef, N., Purdom, E., and Dudoit, S. (2018). Slingshot: Cell lineage and pseudotime inference for single-cell transcriptomics. BMC Genomics 19. 10.1186/s12864-018-4772-0.

26. Saelens, W., Cannoodt, R., Todorov, H., and Saeys, Y. (2019). A comparison of single-cell trajectory inference methods. Nat. Biotechnol. 37, 547–554. 10.1038/s41587-019-0071-9.

27. Farrell, J.A., Wang, Y., Riesenfeld, S.J., Shekhar, K., Regev, A., and Schier, A.F. (2018). Single-cell reconstruction of developmental trajectories during zebrafish embryogenesis. Science (1979). 360. 10.1126/science.aar3131.

28. Dray, N., Than-Trong, E., and Bally-Cuif, L. (2021). Neural stem cell pools in the vertebrate adult brain: Homeostasis from cell-autonomous decisions or community rules? BioEssays 43. 10.1002/bies.202000228.

29. Urbán, N., Blomfield, I.M., and Guillemot, F. (2019). Quiescence of Adult Mammalian Neural Stem Cells: A Highly Regulated Rest. Preprint at Cell Press, 10.1016/j.neuron.2019.09.026 https://doi.org/10.1016/j.neuron.2019.09.026.

30. Labusch, M., Thetiot, M., Than-Trong, E., Morizet, D., Coolen, M., Varet, H., Legendre, R., Ortica, S., Mancini, L., and Bally-Cuif, L. (2024). Prosaposin maintains adult neural stem cells in a state associated with deep quiescence. Stem Cell Reports 19, 515–528. 10.1016/j.stemcr.2024.02.007.

31. Suh, H., Consiglio, A., Ray, J., Sawai, T., D’Amour, K.A., and Gage, F.H.H. (2007). In Vivo Fate Analysis Reveals the Multipotent and Self-Renewal Capacities of Sox2+ Neural Stem Cells in the Adult Hippocampus. Cell Stem Cell 1, 515–528. 10.1016/j.stem.2007.09.002.

32. Letort, G., Foley, T., Mignerey, I., Bally-Cuif, L., and Dray, N. (2025). FishFeats: streamlined quantification of multimodal labeling at the single-cell level in 3D tissues. Preprint, 10.1101/2025.09.02.673708 https://doi.org/10.1101/2025.09.02.673708.

33. Zhou, Y., Bond, A.M., Shade, J.E., Zhu, Y., Davis, C. ha O., Wang, X., Su, Y., Yoon, K.J., Phan, A.T., Chen, W.J., et al. (2018). Autocrine Mfge8 Signaling Prevents Developmental Exhaustion of the Adult Neural Stem Cell Pool. Cell Stem Cell 23, 444–452.e4. 10.1016/j.stem.2018.08.005.

34. Tawarayama, H., Yamada, H., Amin, R., Morita-Fujimura, Y., Cooper, H.M., Shinmyo, Y., Kawata, M., Ikawa, S., and Tanaka, H. (2018). Draxin regulates hippocampal neurogenesis in the postnatal dentate gyrus by inhibiting DCC-induced apoptosis. Sci. Rep. 8. 10.1038/s41598-018-19346-6.

35. Tawarayama, H., Yamada, H., Amin, R., Morita-Fujimura, Y., Cooper, H.M., Shinmyo, Y., Tanaka, H., and Ikawa, S. (2020). Draxin-mediated Regulation of Granule Cell Progenitor Differentiation in the Postnatal Hippocampal Dentate Gyrus. Neuroscience 431, 184–192. 10.1016/j.neuroscience.2020.02.005.

36. Grupp, L., Wolburg, H., and Mack, A.F. (2010). Astroglial structures in the zebrafish brain. Journal of Comparative Neurology 518, 4277–4287. 10.1002/cne.22481.

37. Madelaine, R., and Blader, P. (2011). A cluster of non-redundant Ngn1 binding sites is required for regulation of deltaA expression in zebrafish. Dev. Biol. 350, 198–207. 10.1016/j.ydbio.2010.10.025.

38. Encinas, J.M., Michurina, T. V., Peunova, N., Park, J.H., Tordo, J., Peterson, D.A., Fishell, G., Koulakov, A., and Enikolopov, G. (2011). Division-coupled astrocytic differentiation and age-related depletion of neural stem cells in the adult hippocampus. Cell Stem Cell 8, 566–579. 10.1016/j.stem.2011.03.010.

39. Lupperger, V., Marr, C., and Chapouton, P. (2020). Reoccurring neural stem cell divisions in the adult zebrafish telencephalon are sufficient for the emergence of aggregated spatiotemporal patterns. PLoS Biol. 18. 10.1371/journal.pbio.3000708.

40. Ponti, G., Obernier, K., Guinto, C., Jose, L., Bonfanti, L., and Alvarez-Buylla, A. (2013). Cell cycle and lineage progression of neural progenitors in the ventricular-subventricular zones of adult mice. Proc. Natl. Acad. Sci. U. S. A. 110. 10.1073/pnas.1219563110.

41. Roccio, M., Schmitter, D., Knobloch, M., Okawa, Y., Sage, D., and Lutolf, M.P. (2013). Predicting stem cell fate changes by differential cell cycle progression patterns. Development (Cambridge) 140, 459–470. 10.1242/dev.086215.

42. Kroehne, V., Freudenreich, D., Hans, S., Kaslin, J., and Brand, M. (2011). Regeneration of the adult zebrafish brain from neurogenic radial glia-type progenitors. Development 138, 4831–4841. 10.1242/dev.072587.

43. Grandel, H., Kaslin, J., Ganz, J., Wenzel, I., and Brand, M. (2006). Neural stem cells and neurogenesis in the adult zebrafish brain: Origin, proliferation dynamics, migration and cell fate. Dev. Biol. 295, 263–277. 10.1016/j.ydbio.2006.03.040.

44. Furlan, G., Cuccioli, V., Vuillemin, N., Dirian, L., Muntasell, A.J., Coolen, M., Dray, N., Bedu, S., Houart, C., Beaurepaire, E., et al. (2017). Life-Long Neurogenic Activity of Individual Neural Stem Cells and Continuous Growth Establish an Outside-In Architecture in the Teleost Pallium. Current Biology 27, 3288–3301.e3. 10.1016/j.cub.2017.09.052.

45. Andersen, J., Urbán, N., Achimastou, A., Ito, A., Simic, M., Ullom, K., Martynoga, B., Lebel, M., Göritz, C., Frisén, J., et al. (2014). A Transcriptional Mechanism Integrating Inputs from Extracellular Signals to Activate Hippocampal Stem Cells. Neuron 83, 1085–1097. 10.1016/j.neuron.2014.08.004.

46. Kawaguchi, D., Furutachi, S., Kawai, H., Hozumi, K., and Gotoh, Y. (2013). Dll1 maintains quiescence of adult neural stem cells and segregates asymmetrically during mitosis. Nat. Commun. 4. 10.1038/ncomms2895.

47. Baser, A., Skabkin, M., Kleber, S., Dang, Y., Gülcüler Balta, G.S., Kalamakis, G., Göpferich, M., Ibañez, D.C., Schefzik, R., Lopez, A.S., et al. (2019). Onset of differentiation is post-transcriptionally controlled in adult neural stem cells. Nature 566, 100–104. 10.1038/s41586-019-0888-x.

48. Morris, G.F., and Mathews, M.B. (1989). THE JOURNAL OF BIOLOGICAL CHEMISTRY Regulation of Proliferating Cell Nuclear Antigen during the Cell Cycle*.

49. Kurki,’, P., Vanderlaan, M., Dolbeare, F., Gray, J., Tan’ ’, E.M., and Keck, W.M. (1986). Expression of Proliferating Cell Nuclear Antigen (PCNA)/Cyclin during the Cell Cycle*.

50. Bray, S.J. (2006). Notch signalling: A simple pathway becomes complex. Preprint, 10.1038/nrm2009 https://doi.org/10.1038/nrm2009.

51. de Celis, J.F., and Bray, S. (1997). Feed-back mechanisms affecting Notch activation at the dorsoventral boundary in the Drosophila wing. Development, 3241–3251. 10.1242/dev.124.8.1485.

52. Klein, T., Brennan, K., and Arias, A.M. (1997). An Intrinsic Dominant Negative Activity of Serrate That Is Modulated during Wing Development in Drosophila.

53. Micchelli, C.A., Rulifson, E.J., and Blair, S.S. (1997). The function and regulation of cutexpression on the wing margin of Drosophila: Notch, Wingless and a dominant negative role for Delta and Serrate. Development, 1485–1495. 10.1242/dev.124.8.1485.

54. Miller, A.C., Lyons, E.L., and Herman, T.G. (2009). cis-Inhibition of Notch by Endogenous Delta Biases the Outcome of Lateral Inhibition. Current Biology 19, 1378–1383. 10.1016/j.cub.2009.06.042.

55. Sprinzak, D., Lakhanpal, A., Lebon, L., Santat, L.A., Fontes, M.E., Anderson, G.A., Garcia-Ojalvo, J., and Elowitz, M.B. (2010). Cis-interactions between Notch and Delta generate mutually exclusive signalling states. Nature 465, 86–90. 10.1038/nature08959.

56. Blomfield, I.M., Rocamonde, B., del Mar Masdeu, M., Mulugeta, E., Vaga, S., van den Berg, D.L., Huillard, E., Guillemot, F., and Urbán, N. (2019). Id4 promotes the elimination of the pro-activation factor Ascl1 to maintain quiescence of adult hippocampal stem cells. Preprint, 10.7554/eLife.48561.001 https://doi.org/10.7554/eLife.48561.001.

57. Sueda, R., Imayoshi, I., Harima, Y., and Kageyama, R. (2019). High Hes1 expression and resultant Ascl1 suppression regulate quiescent vs. active neural stem cells in the adult mouse brain. Genes Dev. 33, 511–523. 10.1101/gad.323196.118.

58. Urbán, N., Van Den Berg, D.L.C., Forget, A., Andersen, J., Jeroen, †, Demmers, A.A., Hunt, C., Ayrault, O., and Guillemot, F. Return to quiescence of mouse neural stem cells by degradation of a proactivation protein.

59. Thetiot, M., Taing, L., Morizet, D., Letort, G., and Bally-Cuif, L. (2026). mTORC1 supports progression toward activation competence in quiescent adult neural stem cells. Preprint, 10.64898/2026.05.04.722648 https://doi.org/10.64898/2026.05.04.722648.

60. Coller, H.A., Sang, L., and Roberts, J.M. (2006). A new description of cellular quiescence. PLoS Biol. 4, 0329–0349. 10.1371/journal.pbio.0040083.

61. Fujimaki, K., Li, R., Chen, H., Croce, K. Della, Zhang, H.H., Xing, J., Bai, F., and Yao, G. (2019). Graded regulation of cellular quiescence depth between proliferation and senescence by a lysosomal dimmer switch. Proc. Natl. Acad. Sci. U. S. A. 116, 22624–22634. 10.1073/pnas.1915905116.

62. Kwon, J.S., Everetts, N.J., Wang, X., Wang, W., Della Croce, K., Xing, J., and Yao, G. (2017). Controlling Depth of Cellular Quiescence by an Rb-E2F Network Switch. Cell Rep. 20, 3223–3235. 10.1016/j.celrep.2017.09.007.

63. Pascual-Carreras, E., Garschall, K., and Steinmetz, P.R.H. (2025). Prolonged starvation deepens quiescence in Vasa2/Piwi1-expressing cells of a sea anemone. PLoS Biol. 23, e3003525. 10.1371/journal.pbio.3003525.

64. Rodgers, J.T., King, K.Y., Brett, J.O., Cromie, M.J., Charville, G.W., Maguire, K.K., Brunson, C., Mastey, N., Liu, L., Tsai, C.R., et al. (2014). MTORC1 controls the adaptive transition of quiescent stem cells from G 0 to GAlert. Nature 510, 393–396. 10.1038/nature13255.

65. Bressan, C., Gengatharan, A., Rodriguez-Aller, R., Richter, M.L., Snapyan, M., Fischer-Sternjak, J., Roukerd, M.R., Rosin, N., Cherinet, A., Biernaskie, J., et al. (2026). Cilia beating of ependymal cells regulates adult neural stem cell quiescence via mechanical forces mediated by PKD1/2-TRPM3. Neuron. 10.1016/j.neuron.2026.04.031.

66. Gengatharan, A., Malvaut, S., Marymonchyk, A., Ghareghani, M., Snapyan, M., Fischer-Sternjak, J., Ninkovic, J., Götz, M., and Saghatelyan, A. (2021). Adult neural stem cell activation in mice is regulated by the day/night cycle and intracellular calcium dynamics. Cell 184, 709–722.e13. 10.1016/j.cell.2020.12.026.

67. Marymonchyk, A., Rodriguez-Aller, R., Willis, A., Beaupré, F., Warsi, S., Snapyan, M., Clavet-Fournier, V., Lavoie-Cardinal, F., Kaplan, D.R., Miller, F.D., et al. (2025). Neural stem cell quiescence and activation dynamics are regulated by feedback input from their progeny under homeostatic and regenerative conditions. Cell Stem Cell 32, 445–462.e9. 10.1016/j.stem.2025.01.001.

68. Petrik, D., Myoga, M.H., Grade, S., Gerkau, N.J., Pusch, M., Rose, C.R., Grothe, B., and Götz, M. (2018). Epithelial Sodium Channel Regulates Adult Neural Stem Cell Proliferation in a Flow-Dependent Manner. Cell Stem Cell 22, 865–878.e8. 10.1016/j.stem.2018.04.016.

69. Wang, X., Fujimaki, K., Mitchell, G.C., Kwon, J.S., Della Croce, K., Langsdorf, C., Zhang, H.H., and Yao, G. (2017). Exit from quiescence displays a memory of cell growth and division. Nat. Commun. 8. 10.1038/s41467-017-00367-0.

70. Mitic, N., Neuschulz, A., Spanjaard, B., Schneider, J., Fresmann, N., Novoselc, K.T., Strunk, T., Münster, L., Olivares-Chauvet, P., Ninkovic, J., et al. (2024). Dissecting the spatiotemporal diversity of adult neural stem cells. Mol. Syst. Biol. 20, 321–337. 10.1038/s44320-024-00022-z.

71. Ortica, S., Martinez Herrera, M., Degroux, L., Rochette, B., Dray, N., and Bally-Cuif, L. (2026). Jagged-mediated lateral induction patterns Notch3 signaling within adult neural stem cell populations. Nat. Commun. 17. 10.1038/s41467-026-70478-0.

72. Kassambara, A. (2023). Package “ggcorrplot” Title Visualization of a Correlation Matrix using “ggplot2” ggcorrplot-visualization-of-a-correlation-matrix-using-ggplot2.

73. Robin, X., Turck, N., Hainard, A., Tiberti, N., Lisacek, F., Sanchez, J.C., and Müller, M. (2011). pROC: An open-source package for R and S+ to analyze and compare ROC curves. BMC Bioinformatics 12. 10.1186/1471-2105-12-77.

74. Aigouy, B., Cortes, C., Liu, S., and Prud’Homme, B. (2020). EPySeg: a coding-free solution for automated segmentation of epithelia using deep learning. Development 147, dev194589.

75. Wolf, F.A., Angerer, P., and Theis, F.J. (2018). SCANPY: Large-scale single-cell gene expression data analysis. Genome Biol. 19. 10.1186/s13059-017-1382-0.

76. Virtanen, P., Gommers, R., Oliphant, T.E., Haberland, M., Reddy, T., Cournapeau, D., Burovski, E., Peterson, P., Weckesser, W., Bright, J., et al. (2020). SciPy 1.0: fundamental algorithms for scientific computing in Python. Nat. Methods 17, 261–272. 10.1038/s41592-019-0686-2.

77. Leek, J.T., Johnson, W.E., Parker, H.S., Jaffe, A.E., and Storey, J.D. (2012). The SVA package for removing batch effects and other unwanted variation in high-throughput experiments. Bioinformatics 28, 882–883. 10.1093/bioinformatics/bts034.

78. Johnson, W.E., Li, C., and Rabinovic, A. (2007). Adjusting batch effects in microarray expression data using empirical Bayes methods. Biostatistics 8, 118–127. 10.1093/biostatistics/kxj037.

79. Konopka, T. (2023). umap: Uniform Manifold Approximation and Projection. Preprint.

80. Wickham, H. (2016). HadleyyWickham ggplot2 Elegant Graphics for Data Analysis Second Edition.

81. Kassambara, A. (2026). ggpubr: “ggplot2” Based Publication Ready Plots. Preprint.

82. Paradis, E., and Schliep, K. (2019). Ape 5.0: An environment for modern phylogenetics and evolutionary analyses in R. Bioinformatics 35, 526–528. 10.1093/bioinformatics/bty633.

83. Wickham, H., François, R., Henry, L., Mülle, K., and Vaughan, D. (2026). dplyr: A Grammar of Data Manipulation. Preprint.

84. Lenth, R. V., and Julia Piaskowski, J. (2026). emmeans: Estimated Marginal Means, aka Least-Squares Means. Preprint.

