## Supplementary figures and images for "Linking transcriptome to cell behavior in real time uncovers molecular fate-asymmetry and a quiescence cycle among adult neural stem cells"

### Supplemental Figure S1

Foley et al.
Supplemental Figure S1.

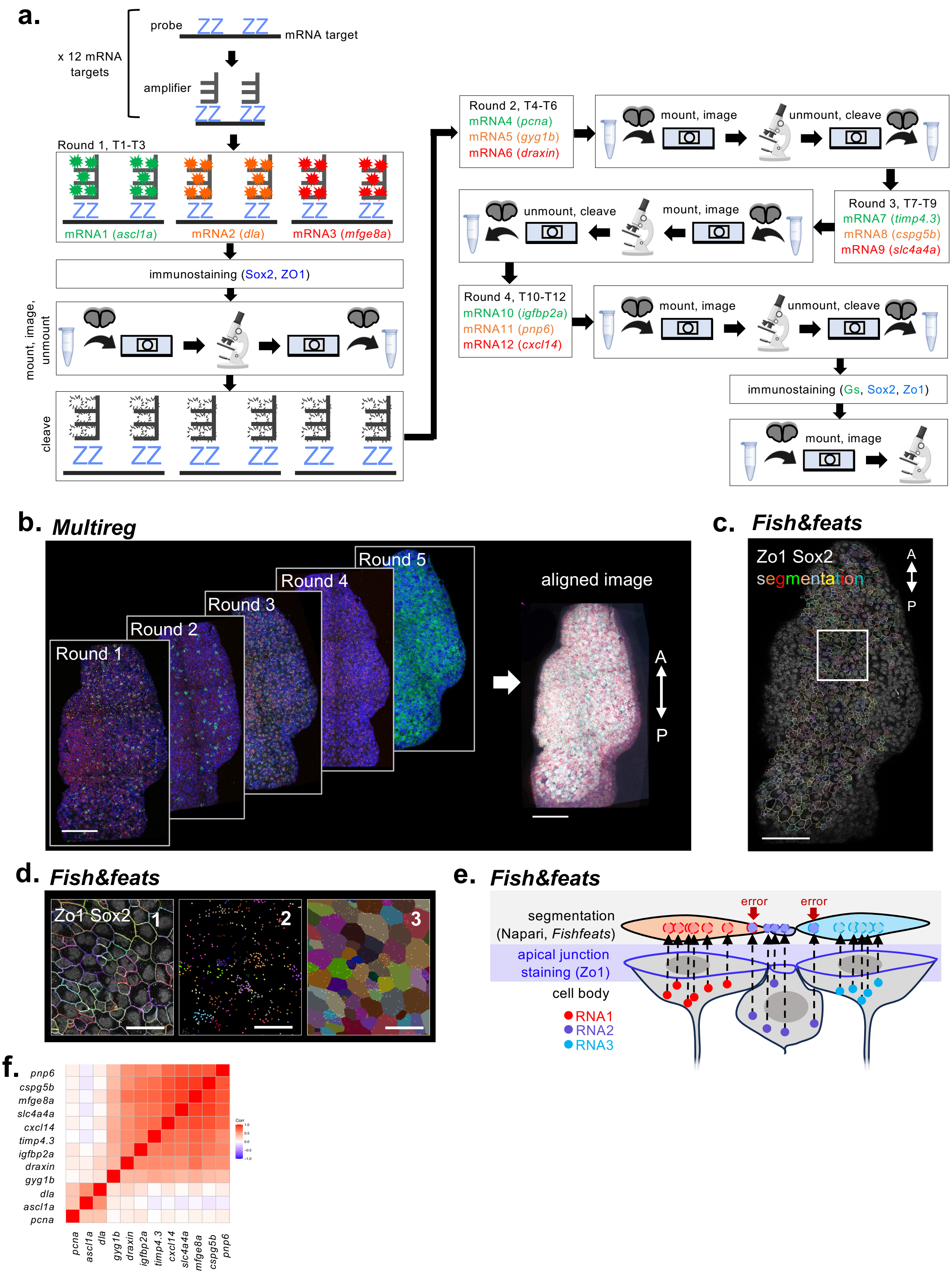

### Supplemental Figure S2

Foley et al.
Supplemental Figure S2.

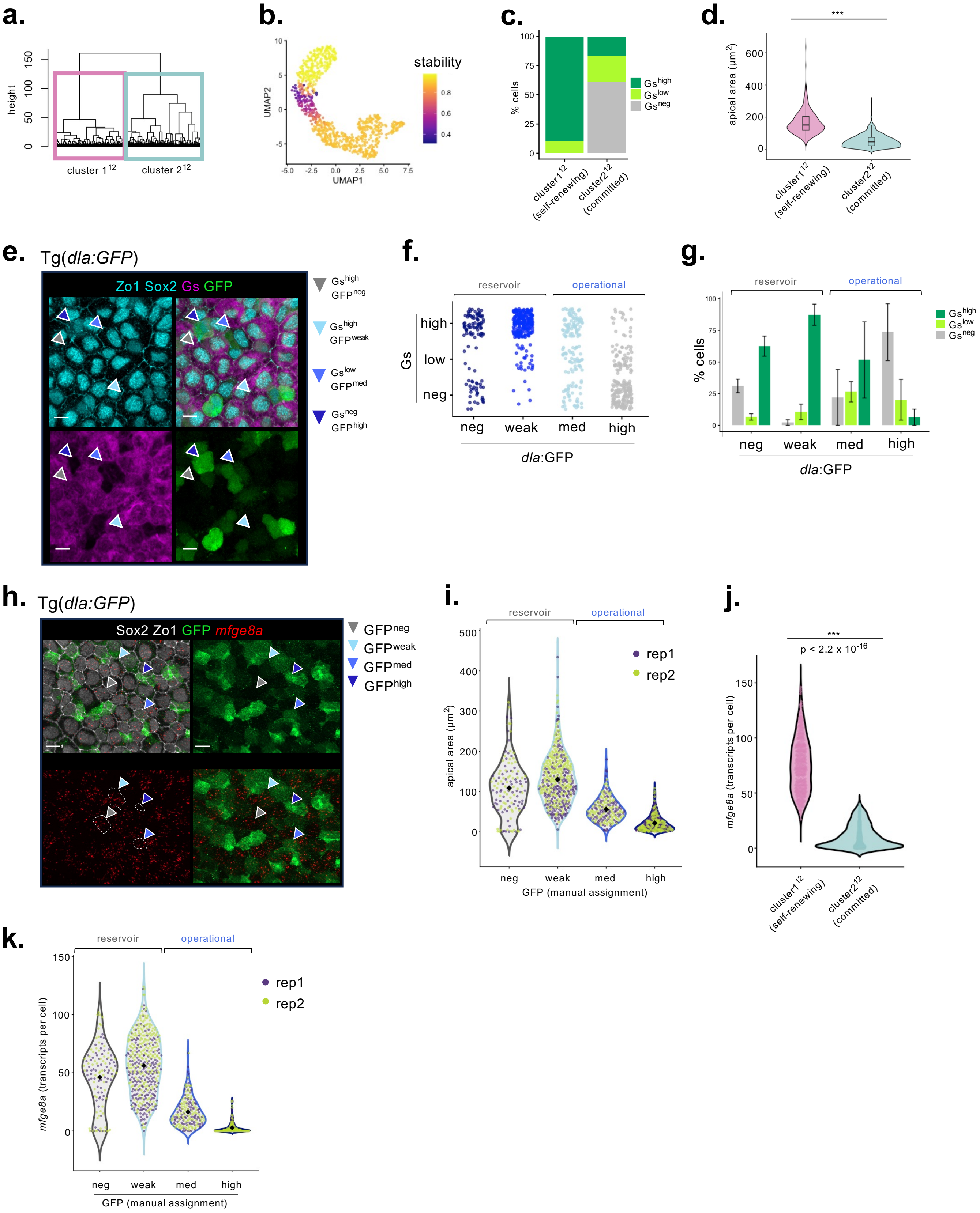

### Supplemental Figure S3

Foley et al.
Supplemental Figure S3.

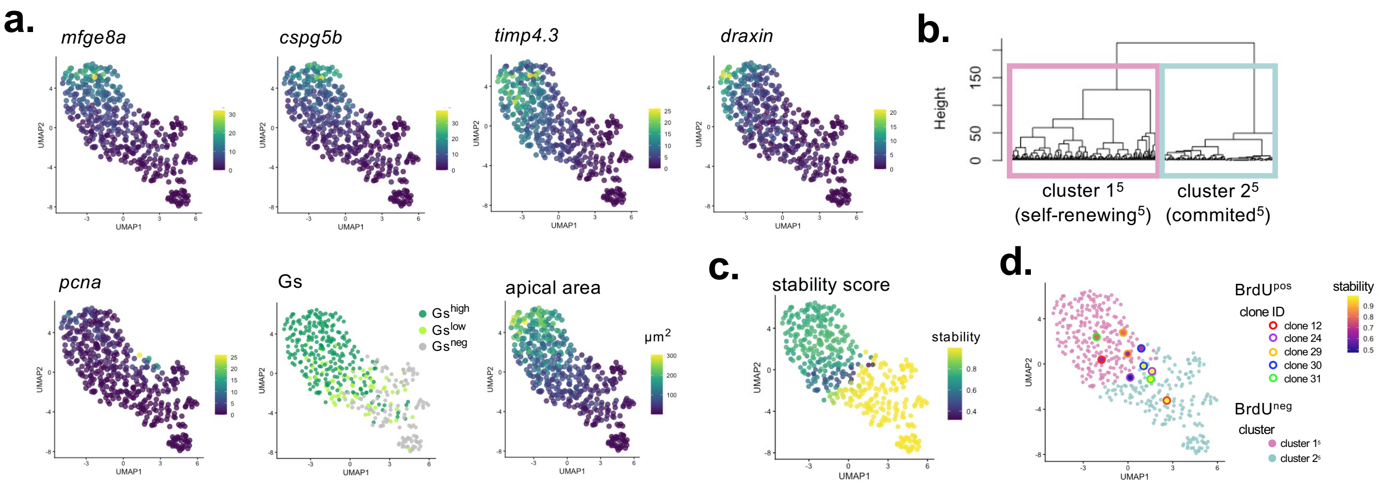

### Supplemental Figure S4

Foley et al.
Supplemental Figure S4.

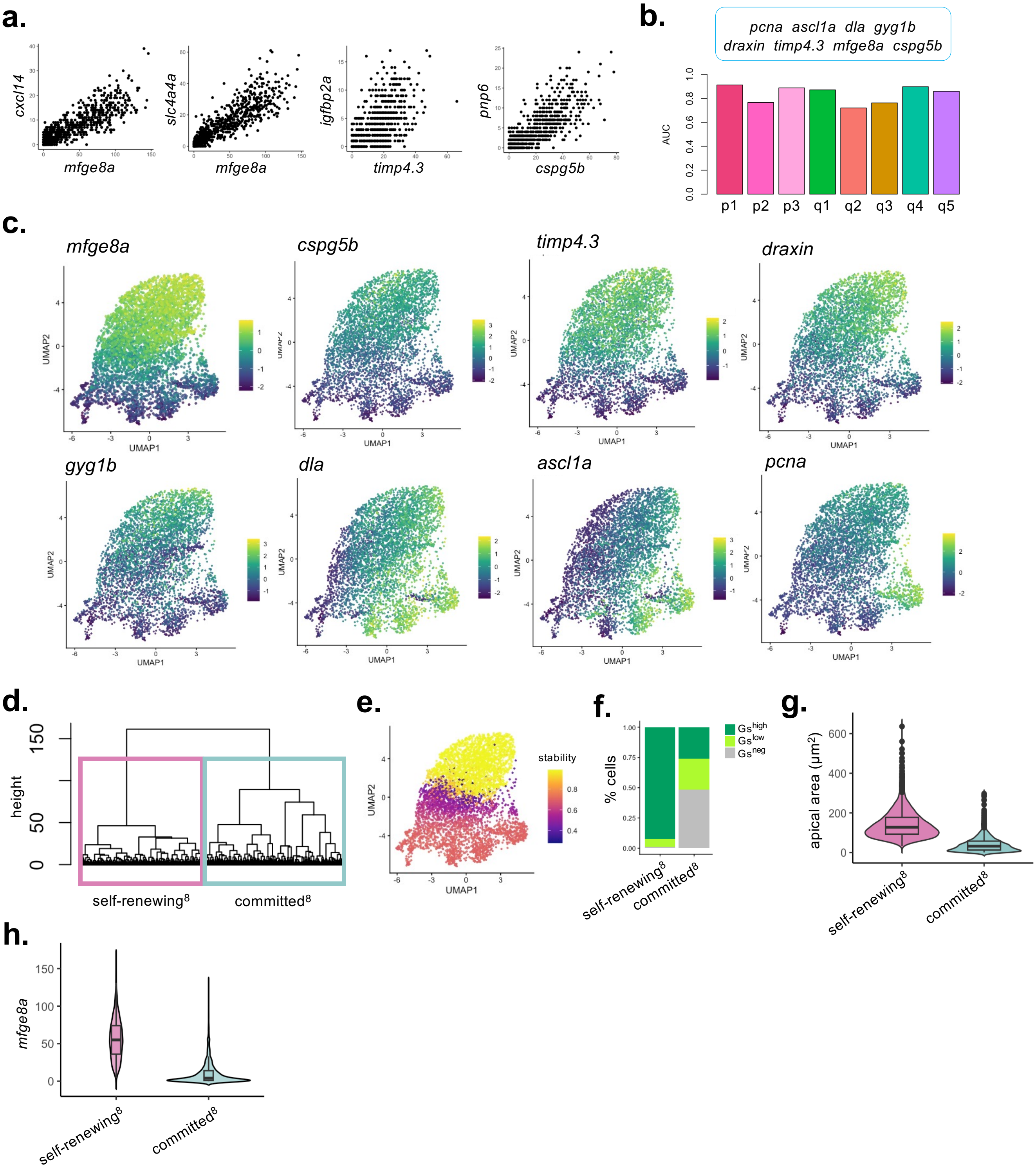

### Supplemental Figure S7

Foley et al.
Supplemental Figure S7.

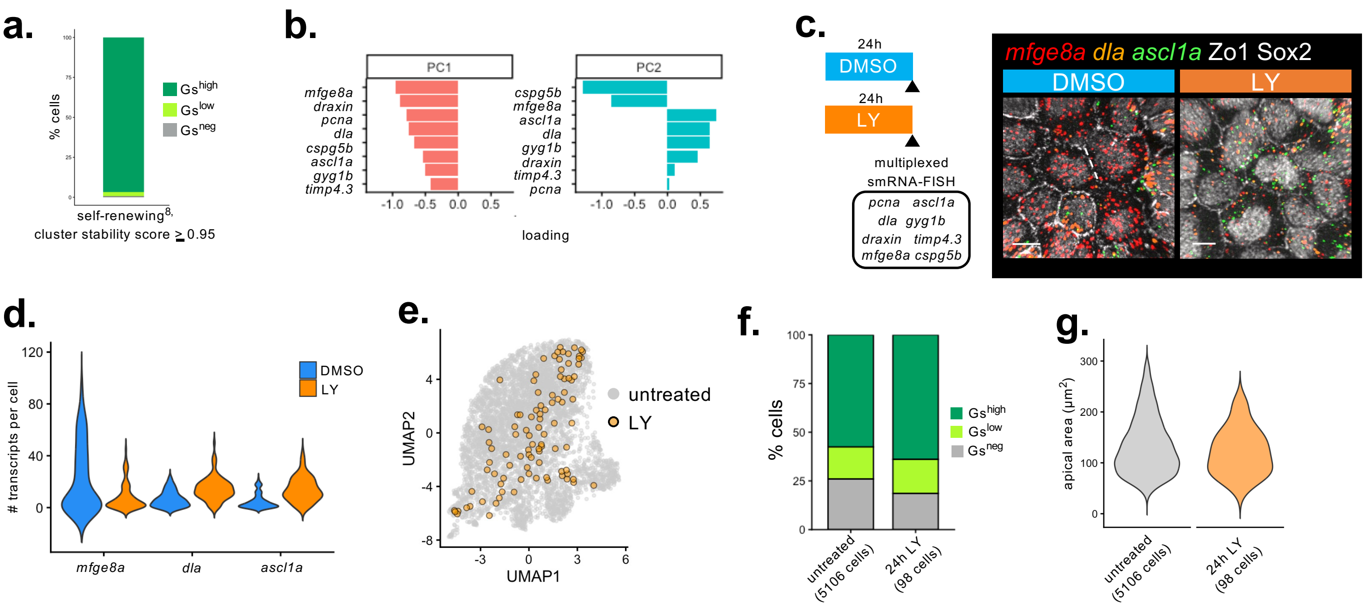
