## Supplemental Figure S5 for "Linking transcriptome to cell behavior in real time uncovers molecular fate-asymmetry and a quiescence cycle among adult neural stem cells"

Foley et al.
Supplemental Figure S5.

**a.**

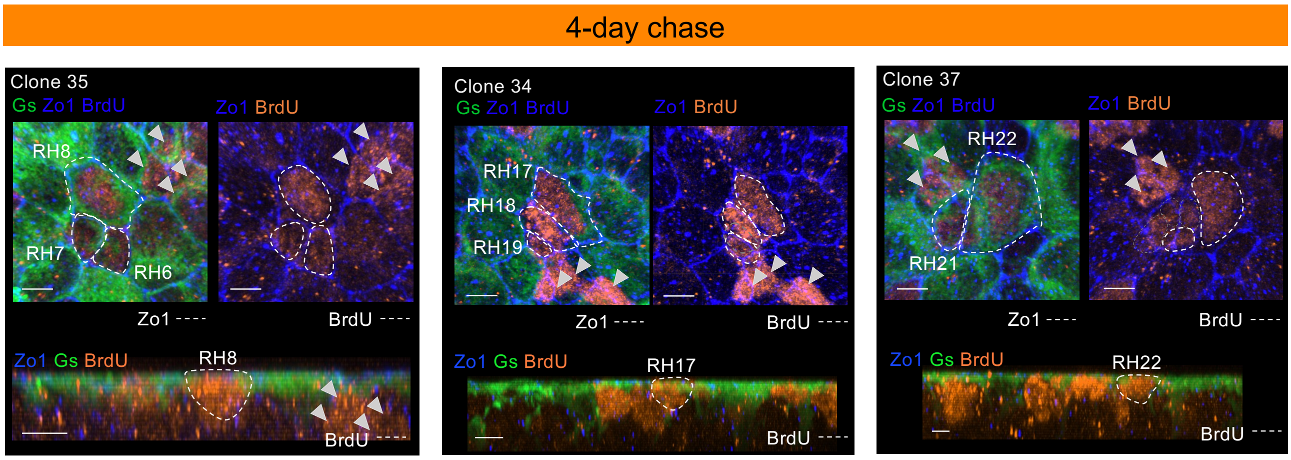


**b.**

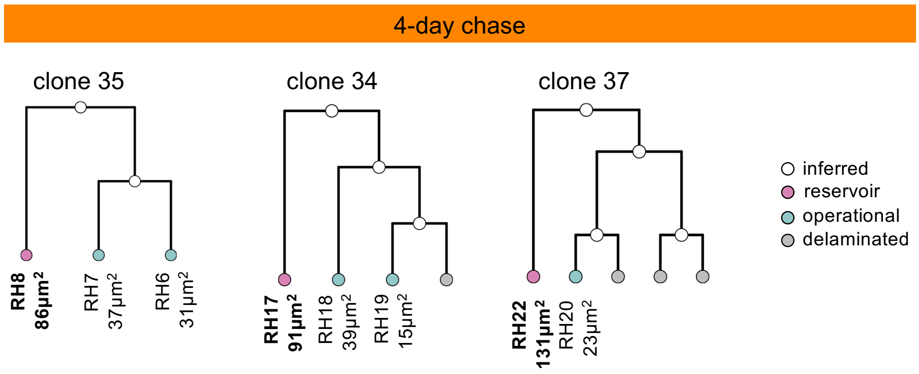

Foley et al.
Supplemental Figure S5.

c.

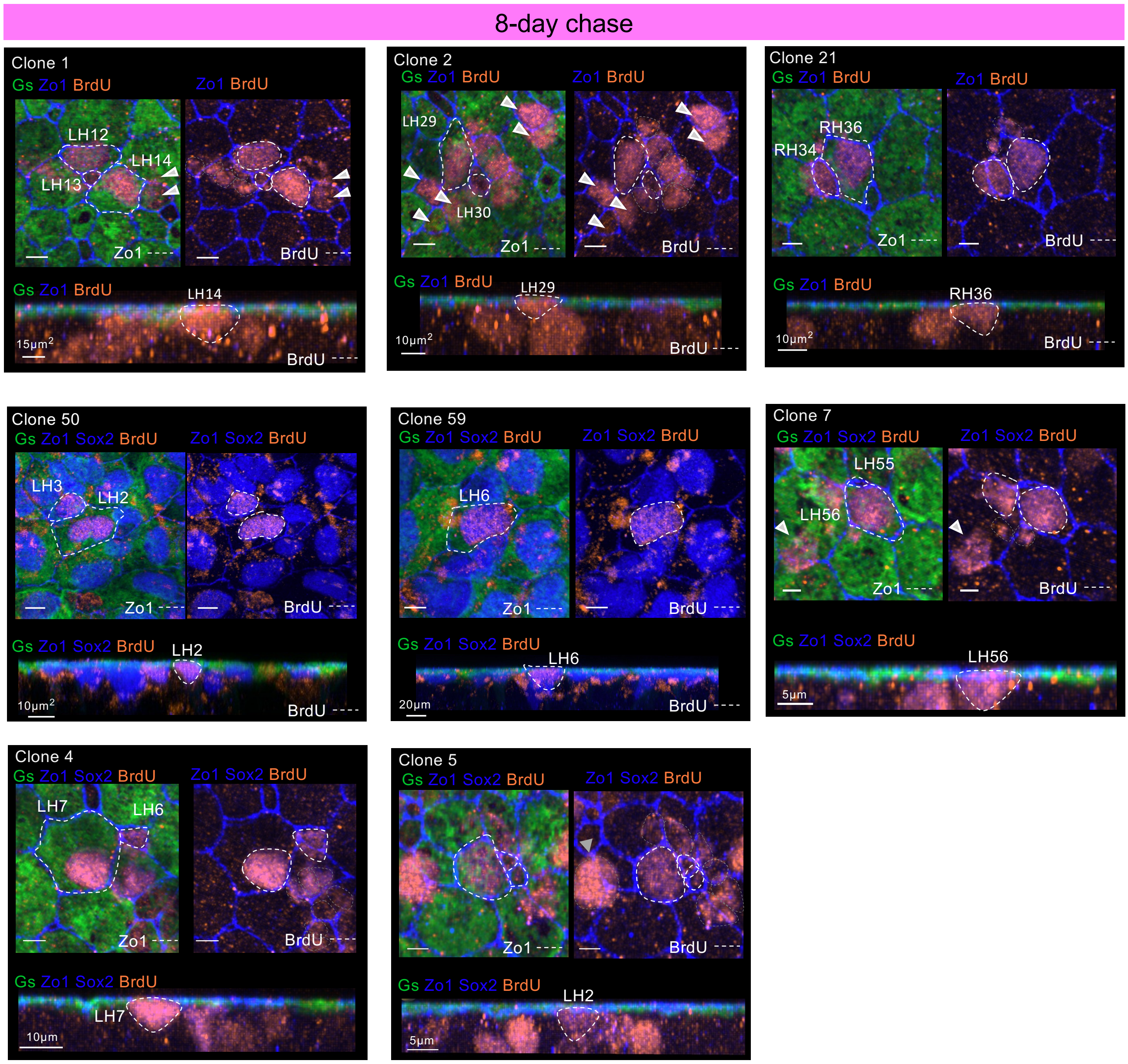


d.

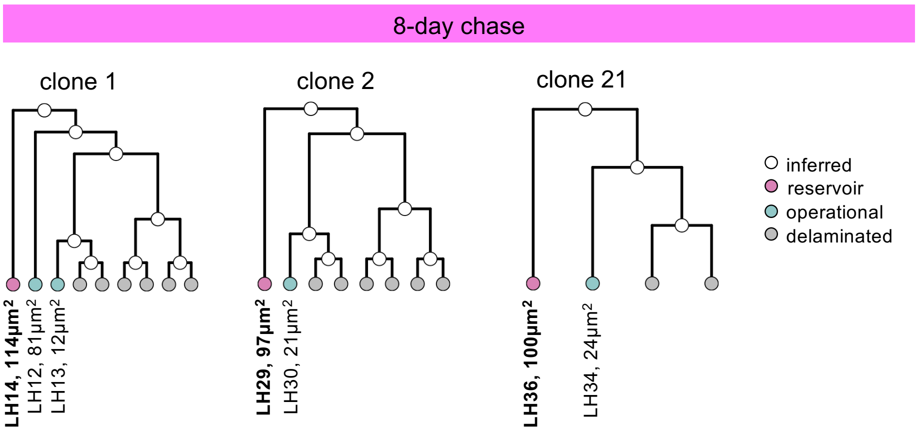

Foley et al.
Supplemental Figure S5. (cont'd)

**e.**

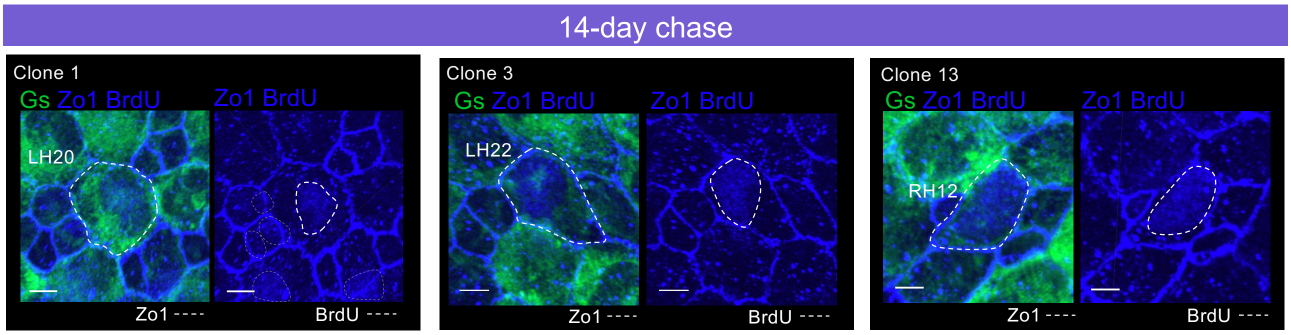


**f.**

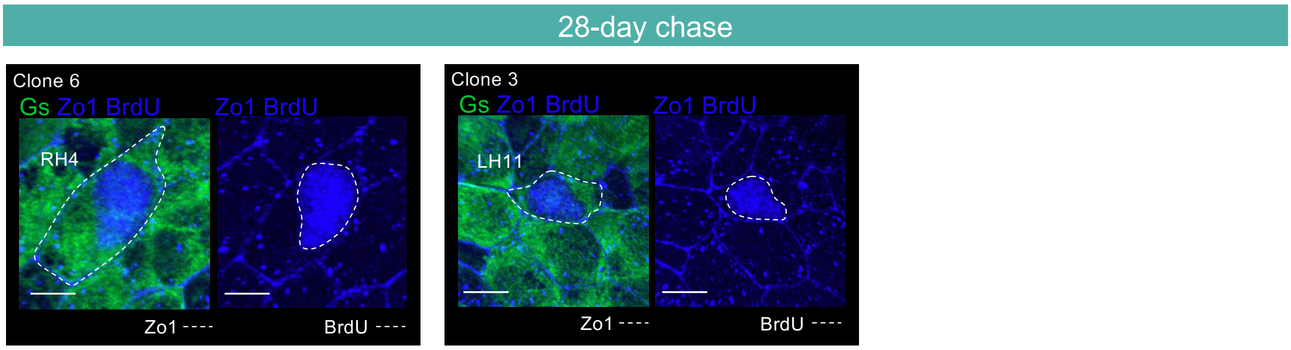
