## Supplemental Figure S6 for "Linking transcriptome to cell behavior in real time uncovers molecular fate-asymmetry and a quiescence cycle among adult neural stem cells"

Foley et al.
Supplemental Figure S6.

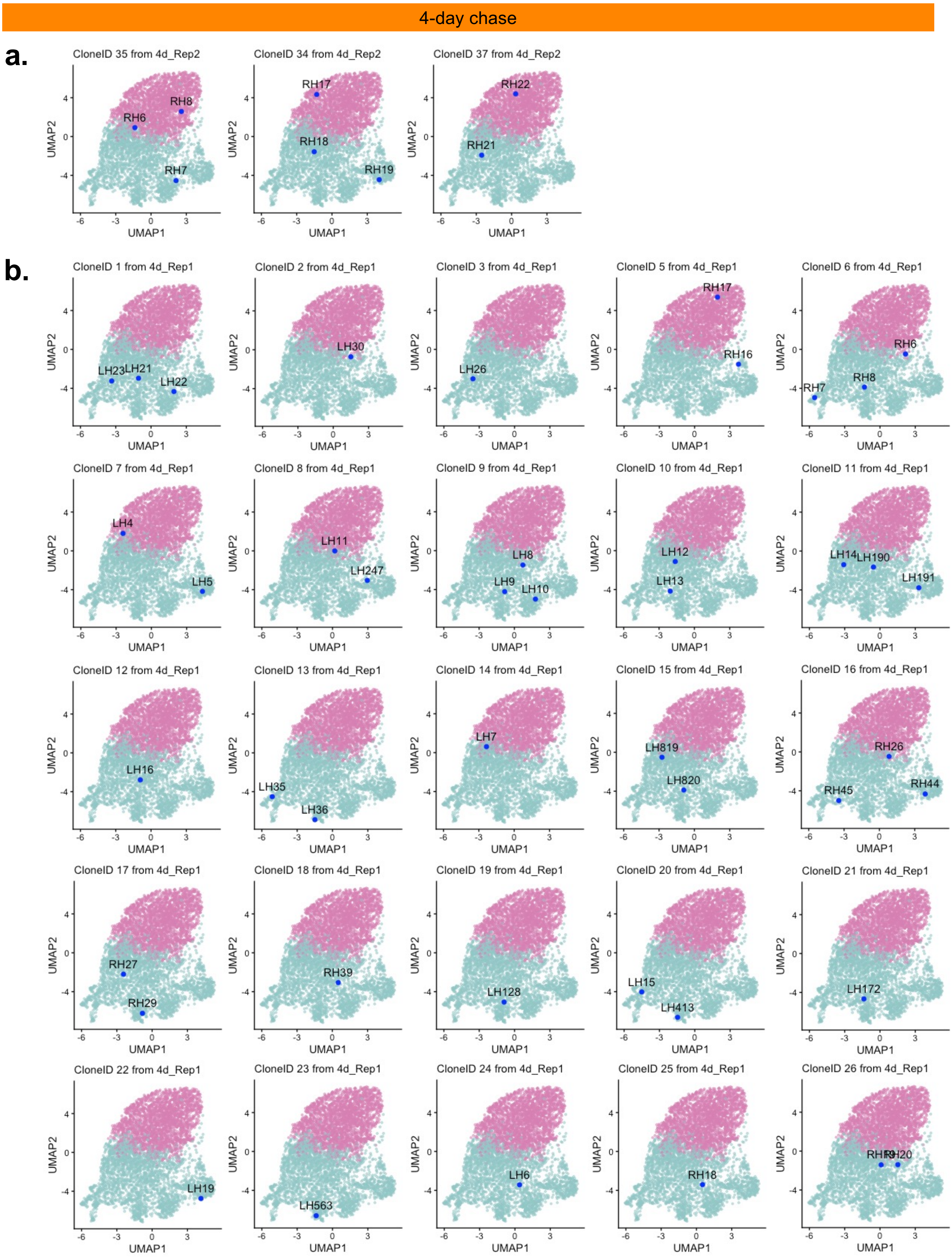

Foley et al.
Supplemental Figure S6 (cont'd).

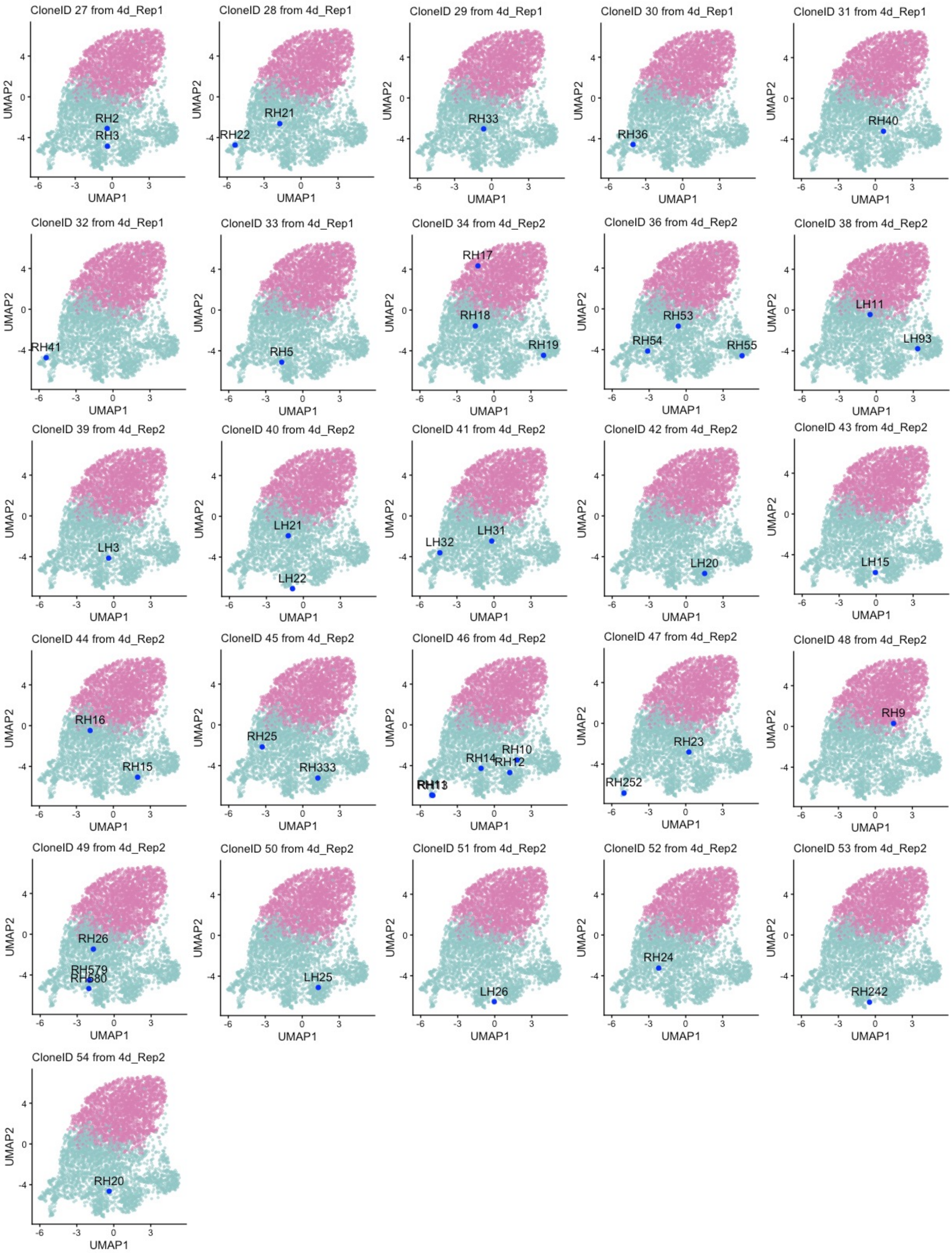

Foley et al.
Supplemental Figure S6 (cont'd).

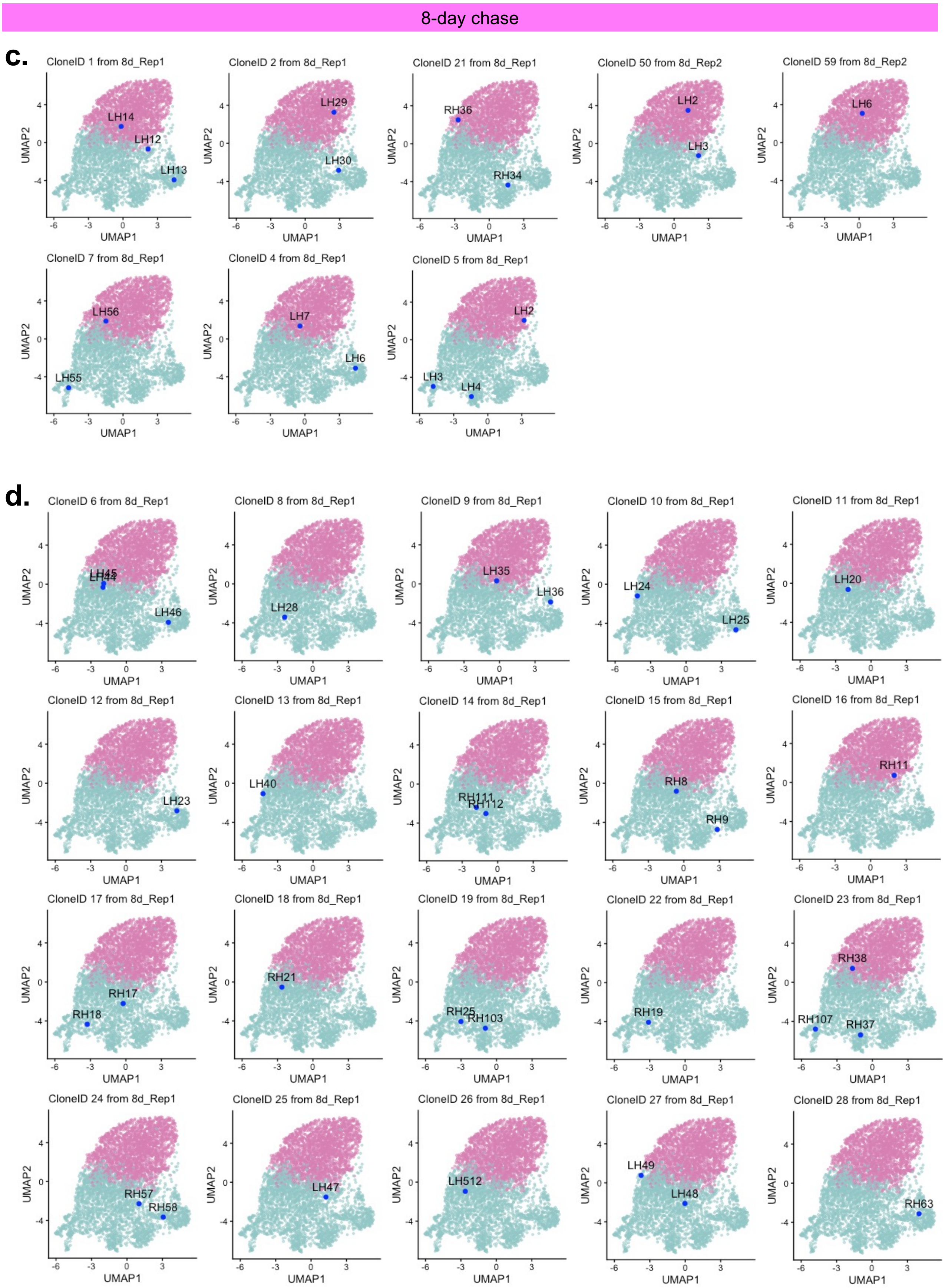

Foley et al.
Supplemental Figure S6 (cont'd).

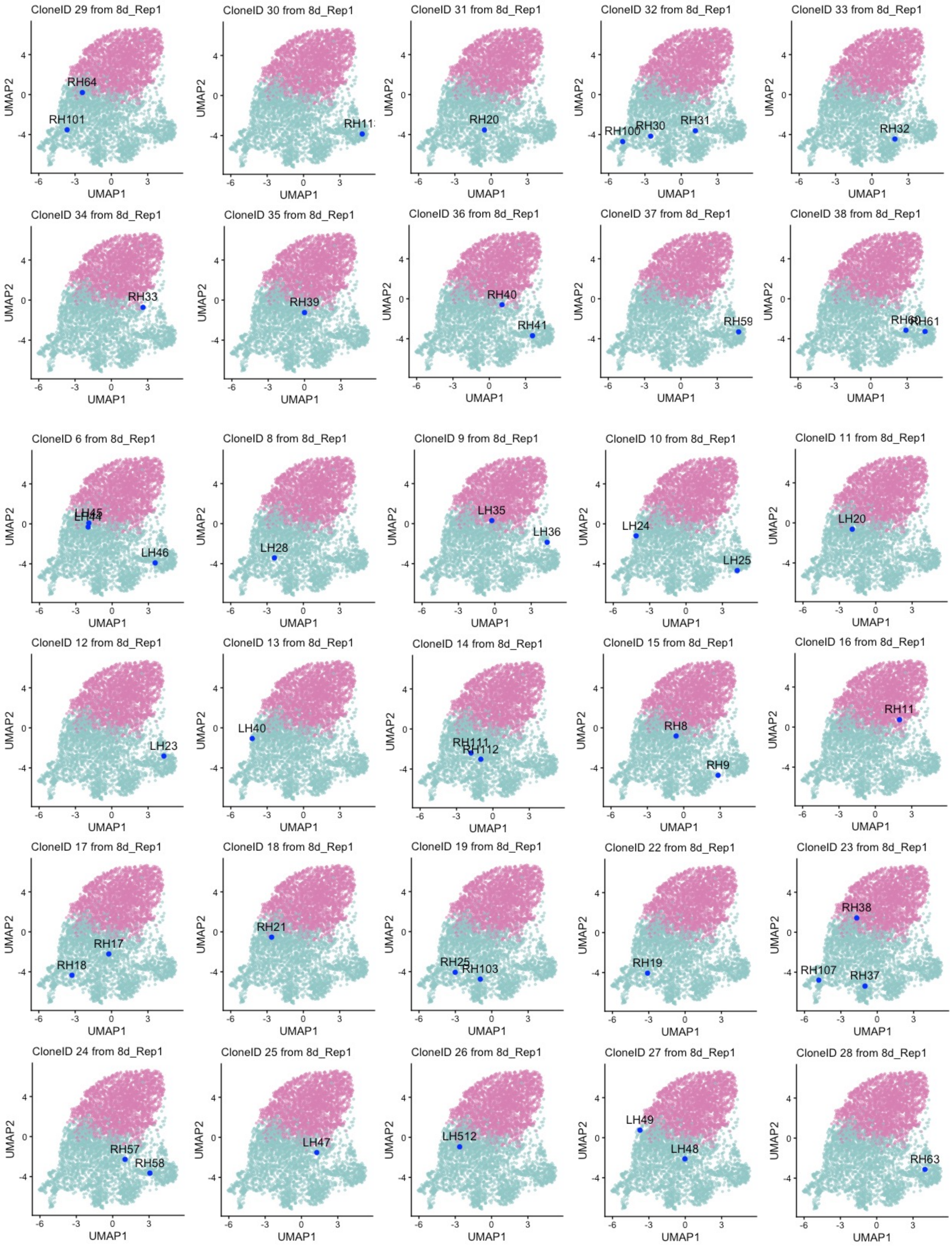

Foley et al.
Supplemental Figure S6 (cont'd).

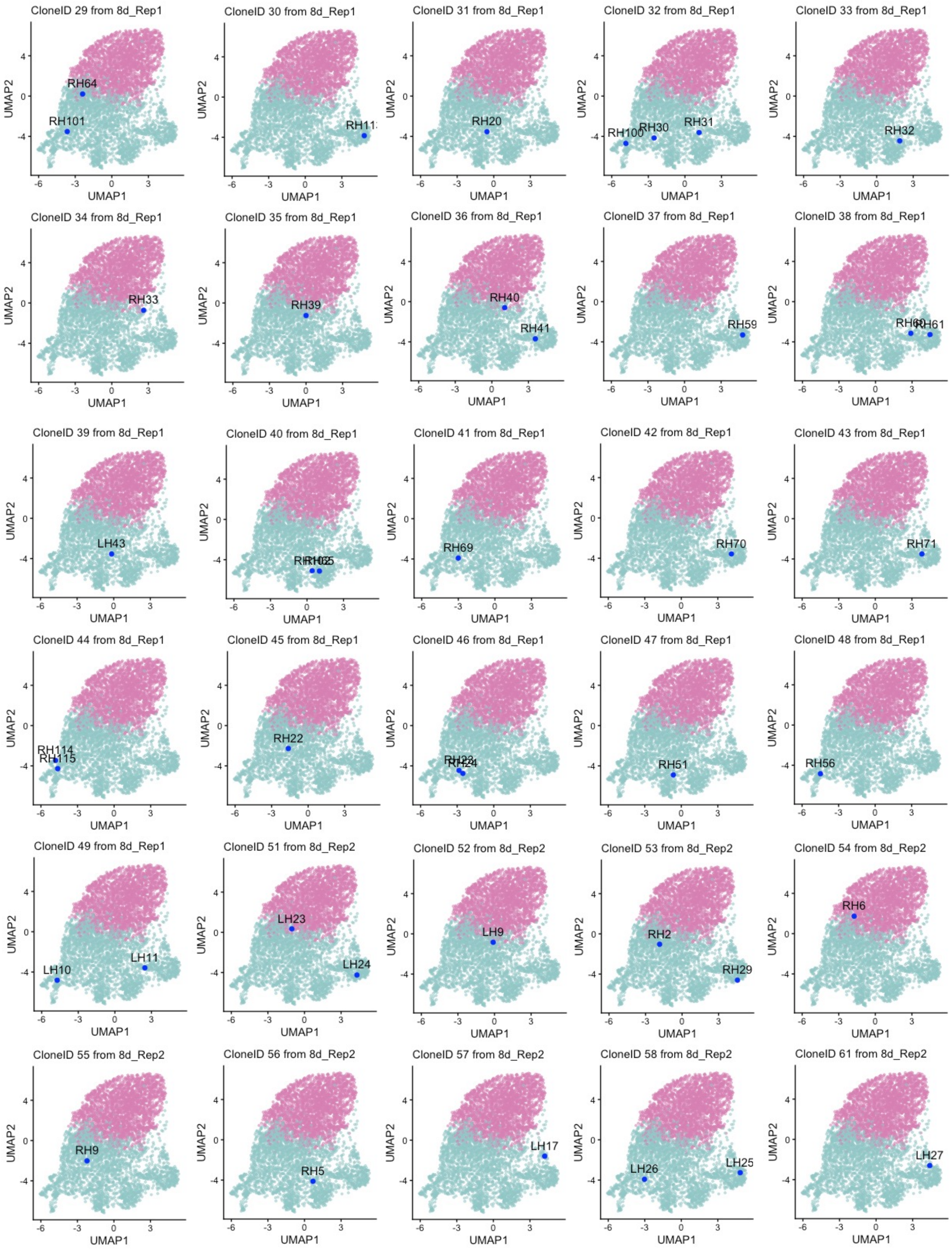

Foley et al.
Supplemental Figure S6 (cont'd).

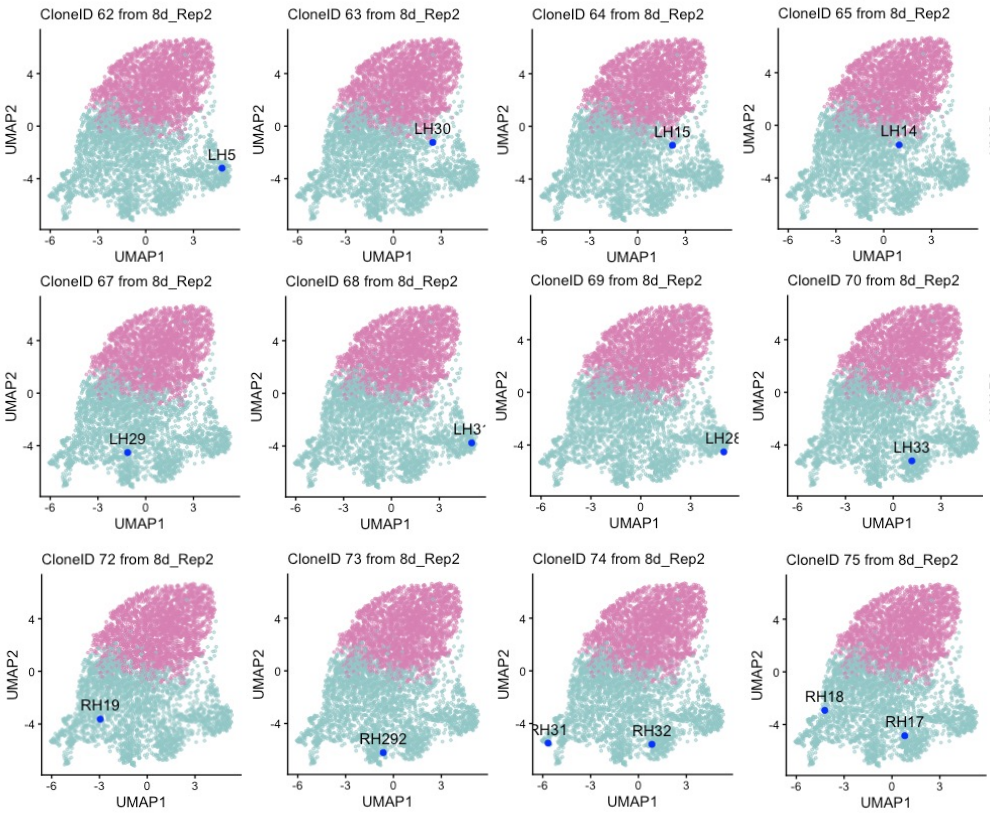


14-day chase

e.

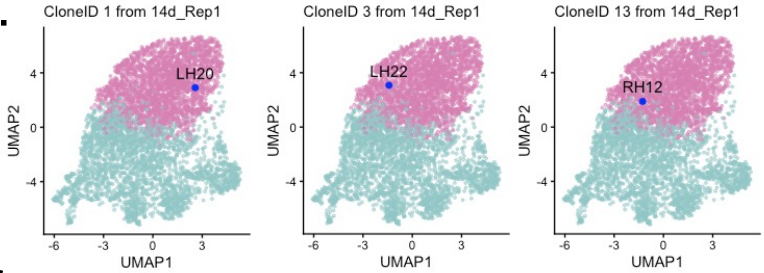


f.

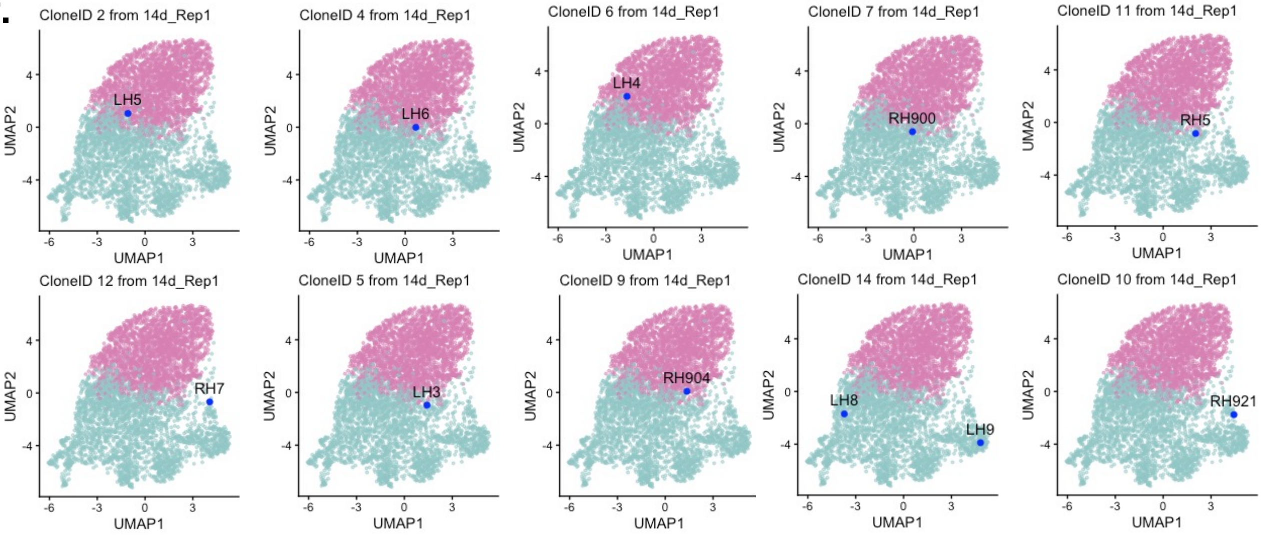

Foley et al.
Supplemental Figure S6 (cont'd).

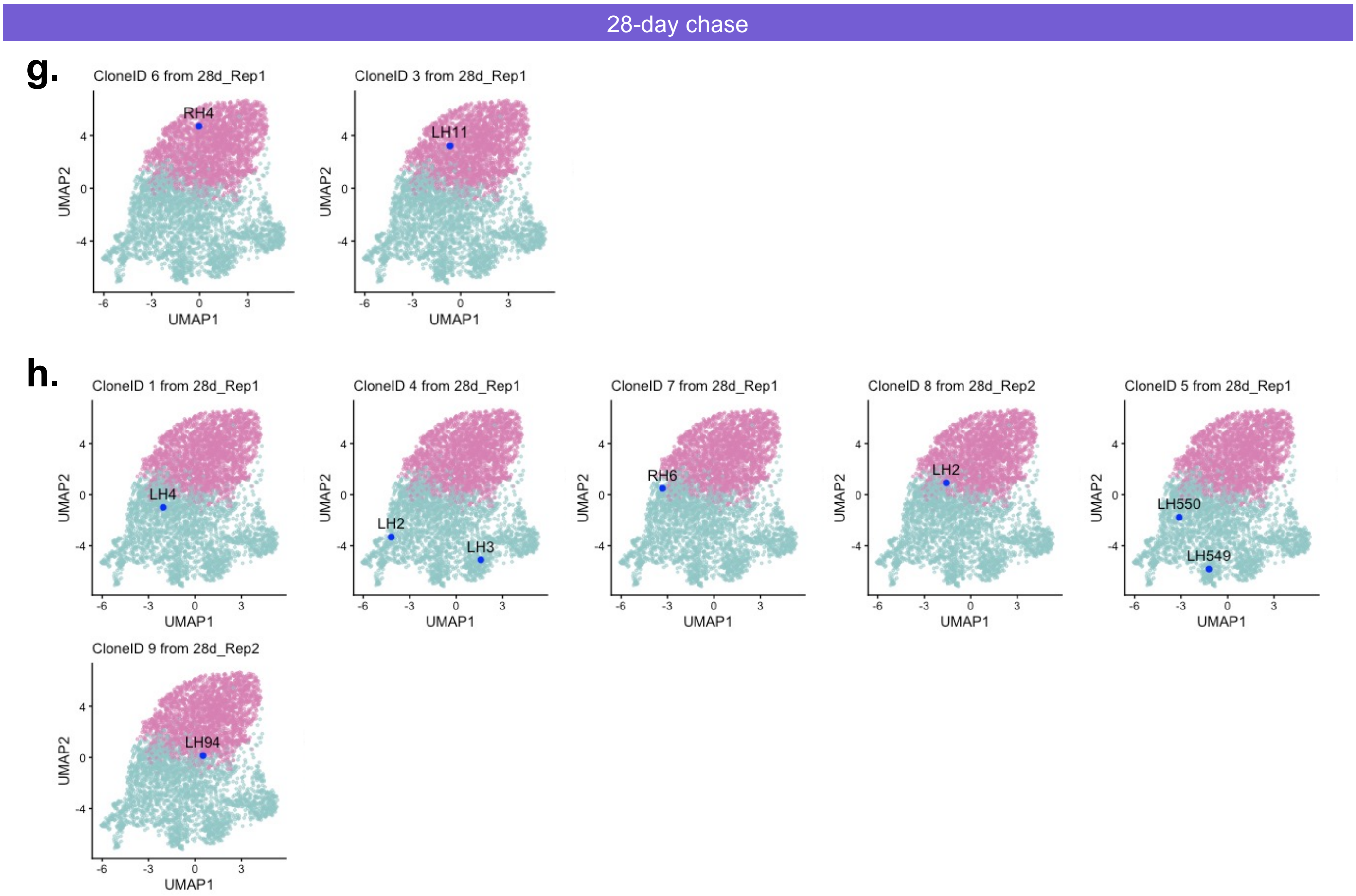
